# *CASTOR1*: A Novel Tumor Suppressor Linking mTORC1 and KRAS Pathways in Tumorigenesis and Resistance to KRAS-Targeted Therapies in Non-Small Cell Lung Cancer

**DOI:** 10.1101/2025.04.23.650349

**Authors:** Xian Wang, Ling Ding, Shenyu Sun, Suet Kee Loo, Luping Chen, Tingting Li, Man-Tzu Wang, Arjun Pennathur, Yufei Huang, Shou-Jiang Gao

## Abstract

Cytosolic arginine sensor for mTORC1 Subunit 1 (CASTOR1) functions as a key regulator of mechanistic target of rapamycin complex 1 (mTORC1) signaling. Despite its frequent dysregulation in cancers via mechanisms such as KSHV microRNA-mediated inhibition or AKT-driven phosphorylation and degradation, the impact of *CASTOR1* loss on tumor initiation and progression remains poorly understood. Here, we identify *CASTOR1* as a critical tumor suppressor in non-small cell lung cancer (NSCLC) by demonstrating that its genetic ablation amplifies tumorigenesis in a *KRAS*-driven genetically engineered mouse model (GEMM;LSL-*KRAS*^G12D^). *CASTOR1* deficiency markedly enhances lung tumor incidence, accelerates tumor progression, and increases proliferative indices in *KRAS^G12D^*-driven tumors (*KRAS^G12D^*;*C1^KO^*) compared to *CASTOR1* wild type (WT) tumors (*KRAS^G12D^*;*C1^WT^*). Advanced-stage tumors exhibit elevated phosphorylated CASTOR1 (pCASTOR1) and reduced total CASTOR1 levels, suggesting active degradation during tumorigenesis. Mechanistically, *CASTOR1* loss amplifies mTORC1 signaling, as evidenced by heightened phosphorylation of downstream effectors 4EBP1 and S6, while also augmenting AKT and ERK activation, uncovering a crosstalk between the PI3K/AKT/mTORC1 and KRAS/ERK pathways. Furthermore, *CASTOR1* ablation induces genome instability, which may contribute to enhanced tumor incidence and progression. Importantly, CASTOR1 deficiency confers resistance to KRAS^G12D^-specific inhibitors, while over half of *KRAS^G12D^*;*C1^WT^* tumors also display resistance. Organoids derived from *KRAS^G12D^*;*C1^KO^* and *KRAS^G12D^*;*C1^WT^* tumors reveal a correlation between KRAS inhibitor resistance and hyperactivation of mTORC1, with mTORC1 and PI3K inhibitors sensitizing resistant tumors to KRAS^G12D^-targeted therapies. These findings position *CASTOR1* as a novel tumor suppressor that modulates mTORC1 and KRAS signaling to constrain NSCLC progression. Our study further highlights the therapeutic potential of combining mTORC1 or ERK inhibitors with KRAS-targeted therapies for NSCLC characterized by hyperactive KRAS signaling and impaired CASTOR1 activity.

**Highlights:**

- *CASTOR1* functions as a tumor suppressor in NSCLC by limiting *KRAS*-driven tumor initiation and progression.
- CASTOR1 is frequently lost or inactivated in wild-type tumors during tumor progression, contributing to advanced-stage malignancies.
- CASTOR1 deficiency amplifies mTORC1 signaling and enhances PI3K/AKT and KRAS/ERK crosstalk, driving tumorigenesis and resistance to KRAS-specific inhibitors.
- Combining mTORC1 or PI3K inhibitors with KRAS-targeted therapies effectively overcomes resistance in *KRAS*-driven NSCLC.

## Introduction

Lung cancer remains the most frequently diagnosed cancer and the leading cause of cancer-related deaths worldwide, accounting for over 1.8 million deaths annually.^1^ Despite significant progress in therapeutic approaches, including immune checkpoint inhibitors and molecularly targeted therapies, the 5-year survival rate for patients with advanced lung cancer remains dismally low, underscoring the need for novel therapeutic approaches.^2^ Non-small cell lung cancer (NSCLC), which comprises 85% of lung cancer cases, is a heterogeneous disease characterized by distinct histological and genomic subtypes, each with unique molecular features and clinical behaviors.^3,4^ Lung adenocarcinoma (LUAD), the most common histological subtype of NSCLC, has been extensively studied, revealing key driver mutations that influence tumor initiation, progression, and therapeutic response.^3,4^

Among all oncogenic driver alterations associated with lung cancer, mutations in the rat sarcoma (*RAS*) gene family are the most prevalent, with Kirsten rat sarcoma (*KRAS*) being the most common subtype.^5,6^ *KRAS* encodes a small GTPase that regulates multiple downstream signaling pathways, including the RAF/MEK/ERK and PI3K/AKT/mTOR cascades, which control cell proliferation, survival, and metabolism.^7^ Oncogenic *KRAS* mutations, often resulting in its constitutive activation, disrupt normal regulatory mechanisms, leading to aberrant signaling, uncontrolled cell growth, and tumor initiation.^7^

*KRAS* gain-of-function mutations are present in approximately 30% of all cancers, with high prevalence in lung cancer, colon cancer, and pancreatic cancer.^7^ In lung cancer, most *KRAS* mutations are missense mutations with substitutions at codons 12 (91%), 13 (5%), or 61 (0.3%). In LUAD, *KRAS* is mutated in approximately 35% of cases.^8^ While significant progress has been made in elucidating the molecular mechanisms underlying *KRAS*-driven oncogenesis, therapeutic targeting of mutant *KRAS* remains challenging, with limited success in the clinic until the recent development of allele-specific inhibitors such as MRTX1133 and sotorasib.^9,10^ Moreover, co-occurring mutations in tumor suppressor genes, such as *TP53*, *STK11*, and *KEAP1*, further complicate therapeutic outcomes and contribute to tumor heterogeneity.^11^

The PI3K/AKT/mTOR pathway is a critical signaling network frequently dysregulated in *KRAS*-driven LUAD. Upregulation of key factors of this pathway such as AKT activity, p-mTORC1, p-S6K, and p-4EBP1 levels are frequently identified in human LUAD patients.^12–15^ mTOR, a serine/threonine kinase, serves as a central regulator of cell growth, metabolism, and autophagy through its two complexes: mechanistic target of rapamycin complex 1 (mTORC1) and complex 2 (mTORC2).^16^ The mTORC1 activity is tightly regulated by nutrient availability, energy status, and growth factor signaling.^16^

Among its regulatory components, cytosolic arginine sensor for mTORC1 subunit 1 (CASTOR1) has emerged as a critical arginine sensor, mediating nutrient-dependent activation of mTORC1.^17^ In the absence of arginine, CASTOR1 dimerizes to interact with the GATOR2 complex (GAP activity towards Rags 2) and inhibits mTORC1 activation. In the presence of arginine, CASTOR1 dissociates from GATOR2, enabling mTORC1 activation. However, high level of CASTOR1 can overcome the inhibitory effect of arginine and suppress mTORC1 activation.^18^ Loss of CASTOR1 leads to hyperactivation of mTORC1, contributing to cancer progression, especially in *KRAS*-driven tumors.^18,19^

Although the role of *CASTOR1* in tumorigenesis remains underexplored, existing studies highlight its importance. CASTOR1 exhibits growth- and tumor-suppressive functions in a KSHV-driven cellular transformation model and a mouse xenograft breast cancer model.^18,20^ In cancer cells with hyperactivated AKT, CASTOR1 is phosphorylated by AKT at S14 (pCASTOR1) and targeted for E3 ligase RNF167-mediated degradation, resulting in arginine-independent mTORC1 activation.^18^ Patients with low *CASTOR1* expression exhibit poorer prognoses across various cancer types, including LUAD.^18,21^ Significantly, a high pCASTOR1 level predicts poorer survival in early-stage male patients with *KRAS* mutations.^19^ In triple-negative breast cancer (TNBC), the downregulator of transcription 1 (DR1) and DR1-associated protein 1 (DRAP1) repressor complex activates mTORC1 by inhibiting CASTOR1 expression, thus promoting tumor progression while conferring mTOR inhibitor everolimus sensitivity.^22^

Given that mTORC1 signaling is often dysregulated in cancer, understanding its regulation in *KRAS*-mutant lung adenocarcinoma is crucial for identifying therapeutic vulnerabilities. In this study, we evaluated the effects of *CASTOR1* loss in a *KRAS^G12D^*-driven genetically engineered mouse (GEM) model of LUAD to test our hypothesis that the loss of *CASTOR1* promotes tumorigenesis through mTORC1 pathway activation. We further investigated the clinical significance of *CASTOR1* in the context of *KRAS*-driven cancer given the historical challenge of targeting KRAS.^9,10^ While simultaneous targeting of mTORC1 and KRAS has shown synergistic effects in NSCLC,^23^ how tumors with hyperactivated mTORC1 caused by low or absent CASTOR1 respond to mTORC1 or KRAS inhibitors remains unknown. Simultaneous targeting of KRAS and other signaling molecules also remain underexplored. We evaluated the effects of mono- and combination therapies involving mTORC1 inhibitor Rapamycin and KRAS^G12D^ inhibitor MRTX1133 by establishing multiple lines of lung cancer organoids derived from *KRAS^G12D^* mouse lung tumors, including both CASTOR1 wild-type (*C1^WT^*) background (*KRAS^G12D^;C1^WT^*) and *CASTOR1* knockout (*C1^KO^*) background (*KRAS^G12D^;C1^KO^*). Notably, we found a high proportion of organoid lines resistant to MRTX1133, particularly in *KRAS^G12D^;C1^KO^* lines but treatment with mTORC1 or PI3K inhibitors sensitized these organoids to MRTX1133.

## Results

### Loss of *CASTOR1* promotes *KRAS^G12D^*-induced cellular transformation

Although the role of *CASTOR1* in tumorigenesis remains largely unexplored, existing studies suggest its tumor-suppressive function in KSHV-induced cancers and breast cancer.^18,20^ Moreover, decreased *CASTOR1* expression is correlated with poorer survival outcomes in at least 10 types of cancer.^18^ To investigate the role of *CASTOR1* in cellular transformation, we generated a *CASTOR1* knockout (*C1^KO^*) GEM model (**Figure S1A**). The successful deletion of *CASTOR1* was confirmed through genotyping PCR, quantitative RT-PCR, and immunoblotting (**Figure S1B-D and 1A**).

We first examined whether *CASTOR1* loss alone could induce cellular transformation. Compared with *C1^WT^* MEFs, which displayed contact inhibition, *C1^KO^* MEFs exhibited morphological changes characterized by overlapping growth patterns and formed small foci in culture (**Figure 1B**). In a soft agar assay, *C1^KO^* MEFs generated small colonies (**Figure 1D**). However, the inability of *C1^KO^* MEFs to form any large foci or robust colonies indicates that loss of *CASTOR1* alone is insufficient to drive full cellular transformation.

**Figure 1.**
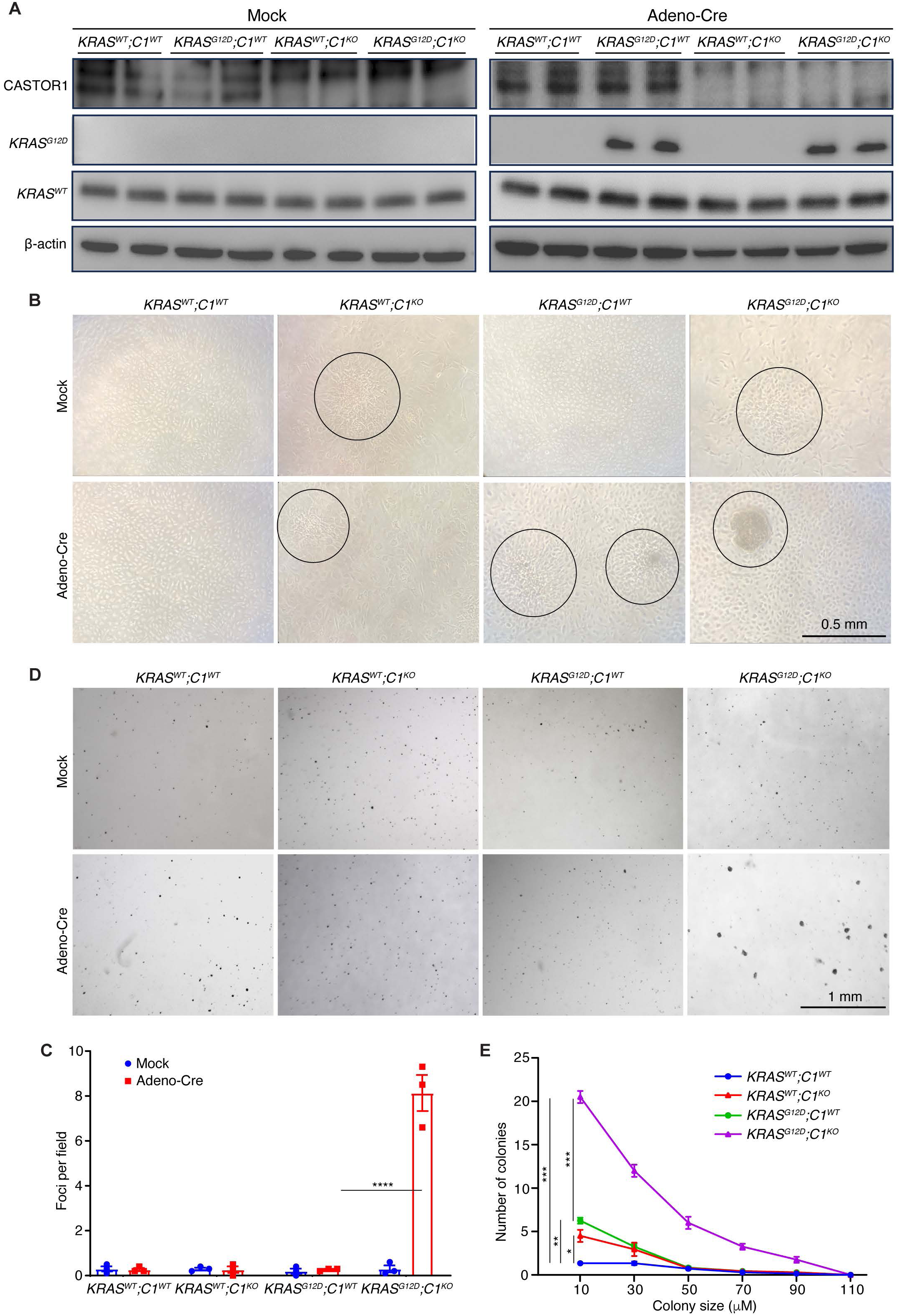
*CASTOR1* ablation enhances *KRAS^G12D^*-driven cellular transformation. (A) Detection of KRAS and KRAS^G12D^ proteins by immunoblotting in passage 2 MEFs isolated from *KRAS^WT^;C1^WT^*, *KRAS^WT^;C1^KO^*, *KRAS^G12D^;C1^WT^* and *KRAS^G12D^;C1^KO^* mice. The expression of *KRAS^G12D^* in MEFs was induced by Adeno-Cre infection and KRAS^G12D^ protein was examined at day 3 post-infection. (B-C) *CASTOR1* ablation enhances *KRAS^G12D^*-induced foci formation in culture. Mock- or Adeno-Cre-infected cells were seeded at 5×10^5^ cells/well in 10 cm-dish at day 3 post-infection and cultured with daily change of medium. Foci formation was examined at day 14 post-seeding with representative images shown (B) and quantified (C). (D-E) *CASTOR1* ablation enhances *KRAS^G12D^*-induced colony formation in semi-soft agar. Mock- or Adeno-Cre-infected cells were seeded in soft agar at day 3 post-infection with daily change of medium. Colonies were examined at day 14 post-seeding with representative images shown (D) and colonies from Adeno-Cre-infected cells were quantified (E). Statistical analysis was performed by Student’s t-test.

Given the high prevalence of *KRAS* mutations in cancer, we next investigated whether *CASTOR1* loss could enhance transformation driven by a mutant *KRAS*. *C1^KO^* mice were crossed with *KRAS^G12D^* mice carrying a stop-floxed *KRAS^G12D^* transgene to generate *KRAS^G12D^*;*C1^KO^* mice (**Figure S2**). Upon infection with Cre recombinase-expressing adenovirus (Adeno-Cre), MEFs expressed *KRAS^G12D^* (**Figure1A and S3**). As expected, *KRAS^G12D^* expression in MEFs induced the formation of small foci in culture (**Figure1B-C**) and small colonies in soft agar (**Figure1D-E**). Notably, *CASTOR1* loss significantly increased the formation of *KRAS^G12D^*-induced foci (**Figure1B-C**). Similarly, *CASTOR1* loss enhanced *KRAS^G12D^*-driven colony formation in soft agar (**Figure1D-E**). These findings indicate that *CASTOR1* loss synergizes with *KRAS^G12D^* to promote cellular transformation.

### Loss of *CASTOR1* enhances *KRAS^G12D^*-driven lung adenocarcinoma incidence and progression

To assess whether *CASTOR1* loss predisposes to cancer in vivo, we monitored *C1^KO^* mice but observed no significant increase in spontaneous tumor formation compared to *C1^WT^* controls (data not shown). However, clinical data show that lower *CASTOR1* expression correlates with worse survival outcomes in LUAD and LUSC,^18^ and elevated pCASTOR1 levels are associated with poor survival in male patients with *KRAS* mutations at early disease stages.^19^ Thus, we examined the impact of *CASTOR1* loss on LUAD incidence and progression using the *KRAS^G12D^* mouse model. Following Adeno-Cre administration for 5 months, both *KRAS^G12D^;C1^WT^* and *KRAS^G12D^;C1^KO^* mice developed lung tumor nodules (**Figure 2A**). Notably, *KRAS^G12D^;C1^KO^* mice exhibited a significant higher number of nodules, larger average nodule size, and greater overall lung lesion burden compared to *KRAS^G12D^;C1^WT^* mice (**Figure 2A**). These results demonstrate that *CASTOR1* loss enhances *KRAS^G12D^*-driven LUAD tumor incidence and accelerates tumor growth.

**Figure 2.**
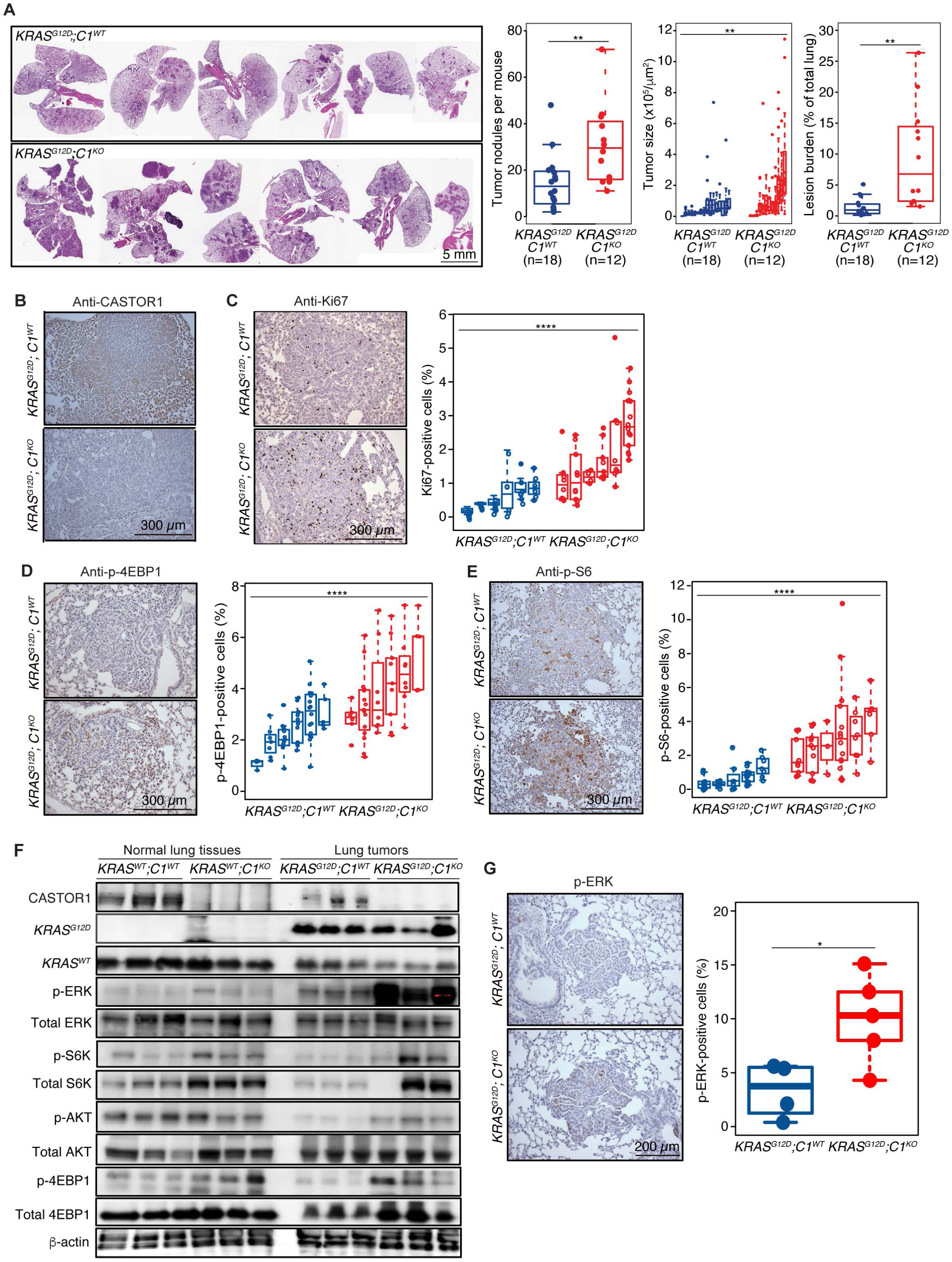
*CASTOR1* has a tumor suppressor function in a *KRAS^G12D^*-driven murine model of lung cancer. (A) Representative H&E images of lungs collected from *KRAS^G12D^;C1^WT^* or *KRAS^G12D^;C1^KO^* mice 5 months after intranasal inoculation with Adeno-Cre (left panel). Tumor incidence, size and burden (tumor area/total lung area ×100) were quantified (left, middle and right panels, respectively). Data presented as mean +/- standard error of the mean (SEM). Statistical analysis was performed by nested-Anova and Student’s t-test. (B) Representative IHC staining of CASTOR1 in lung tumors from *KRAS^G12D^;C1^WT^* or *KRAS^G12D^;C1^KO^* mice. (C) Representative IHC staining images of Ki67 in lung tumors from *KRAS^G12D^;C1^WT^* or *KRAS^G12D^;C1^KO^* mice (left panel) and quantification of Ki67-positive cells (right panel). Statistical analysis was performed by nested-Anova. (D and E) Representative IHC staining images of p-4EBP1 (D) and p-S6 (E) of lung tumors from *KRAS^G12D^;C1^WT^* or *KRAS^G12D^;C1^KO^* mice (left panels) and quantification of positive cells (right panels). Statistical analysis was performed by nested-Anova. (F) Immunoblotting analysis of mTORC1 downstream targets S6K and 4EBP1, KRAS downstream target ERK and AKT, and their phosphorylated forms in normal lung tissues from *KRAS^WT^;C1^WT^* and *KRAS^WT^;C1^KO^* mice, and tumors from *KRAS^G12D^;C1^WT^ and KRAS^G12D^;C1^KO^* mice. Three samples per group were analyzed. (G). Representative IHC staining images of p-ERK in lung tumors from *KRAS^G12D^;C1^WT^* or *KRAS^G12D^;C1^KO^* mice (left panel) and quantification of positive cells (right panel). Statistical analysis was performed by Student’s t-test.

### Ablation of *CASTOR1* enhances mTOR signaling in *KRAS^G12D^*-induced lung adenocarcinoma

To confirm *CASTOR1* ablation in *KRAS^G12D^*-driven LUAD, we examined CASTOR1 expression in tumors and adjacent normal tissues. CASTOR1 expression was evident in *KRAS^G12D^;C1^WT^* tumors but was notably reduced compared to adjacent normal tissues, whereas no CASTOR1 expression was detected in *KRAS^G12D^;C1^KO^* tumors or adjacent tissues (**Figure 2B**). This confirmed the complete loss of *CASTOR1* in *KRAS^G12D^;C1^KO^* tumors.

Next, we assessed tumor proliferation by analyzing Ki67 expression. *KRAS^G12D^;C1^KO^* tumors displayed a significantly higher proliferation index compared to *KRAS^G12D^;C1^WT^* tumors, indicating accelerated tumor growth in the absence of CASTOR1 (**Figure 2C**).

Previous studies have shown that CASTOR1 downregulation leads to activated mTORC1 signaling pathway.^17,18^ mTORC1 activation promotes protein synthesis by phosphorylating downstream targets, such as S6K1 and 4EBP1.^16^ This phosphorylation inhibits the translational repressor function of 4EBP1 and enhances ribosomal activity, driving cell proliferation. To investigate whether increased mTORC1 activity contributed to the elevated proliferation in *KRAS^G12D^;C1^KO^* tumors, we analyzed phosphorylation levels of mTORC1 downstream targets, 4EBP1 and S6. *KRAS^G12D^;C1^KO^* tumors exhibited significantly higher levels of phosphorylated 4EBP1 (p-4EBP1) and phosphorylated S6 (p-S6) compared to *KRAS^G12D^;C1^WT^* tumors (**Figure 2D-E**). These findings demonstrate that *CASTOR1* loss leads to heightened mTORC1 signaling, which likely drives enhanced tumor proliferation and development.

### CASTOR1 ablation promotes lung tumor development by activating both mTORC1 and KRAS/ERK pathways

Previous studies identified CASTOR1 as a negative regulator of the canonical mTORC1 signaling pathway.^17,18^ To further explore its role, we compared the activation of key signaling pathways in normal lung tissues between *KRAS^WT^;C1^WT^* and *KRAS^WT^;C1^KO^* mice, and in tumors between *KRAS^G12D^;C1^WT^* and *KRAS^G12D^;C1^KO^* mice (**Figure 2F**). As expected, *CASTOR1* ablation led to constitutive activation of the mTORC1 pathway, evidenced by elevated levels of p-S6K and p-4EBP1 in normal lung tissues from *KRAS^WT^;C1^KO^* mice compared to *KRAS^WT^;C1^WT^* mice. A similar pattern was observed in *KRAS^G12D^*-induced tumors, confirming the regulatory role of CASTOR1 in mTORC1 signaling.

Interestingly, tumors with *CASTOR1* ablation also displayed increased phosphorylation of AKT at serine 473 (p-AKT-S473), suggesting a potential negative feedback loop between CASTOR1 and the PI3K/AKT pathway. Previous reports indicate that *KRAS^G^*^12^ promotes mTORC1 activation through the MEK/ERK/ROS axis and the PI3K/AKT/mTORC1 pathways.^24^ Consistent with this, we observed elevated levels of p-ERK in both normal lung tissues and tumors lacking CASTOR1. However, the activation of p-ERK was significantly more robust in tumors, implying that heightened ERK signaling may also contribute to tumor development in the absence of CASTOR1. IHC staining further validated these findings, showing markedly elevated p-ERK levels in *KRAS^G12D^;C1^KO^* tumors compared to *KRAS^G12D^;C1^WT^* tumors (**Figure 2G**).

Taken together, *CASTOR1* ablation drives tumor growth by activating the mTORC1 pathway and amplifying *KRAS^G12D^*-induced ERK pathway activation. These dual effects underscore the critical role of CASTOR1 in modulating key oncogenic signaling pathways.

### CASTOR1 is inactivated during lung tumor progression

Our findings thus far highlight a tumor-suppressive role for *CASTOR1*. Examination of TCGA and other cancer databases revealed no detectable association between *CASTOR1* deletions or mutations, including point mutations, and any cancer types (data not shown) despite lower *CASTOR1* expression levels were correlated with poorer prognosis.^18^ These observations suggest that CASTOR1 is inactivated during tumor progression, distinguishing it from classical tumor suppressors, where mutations or deletions are often prerequisites for cancer development.

Previous studies have shown that AKT-mediated phosphorylation of CASTOR1 leads to its degradation.^18^ To determine whether this mechanism might be involved in tumor progression, we analyzed CASTOR1 and pCASTOR1 levels in consecutive tumor sections. A negative correlation between total CASTOR1 and pCASTOR1 levels was observed (**Figure 3A**).

**Figure 3.**
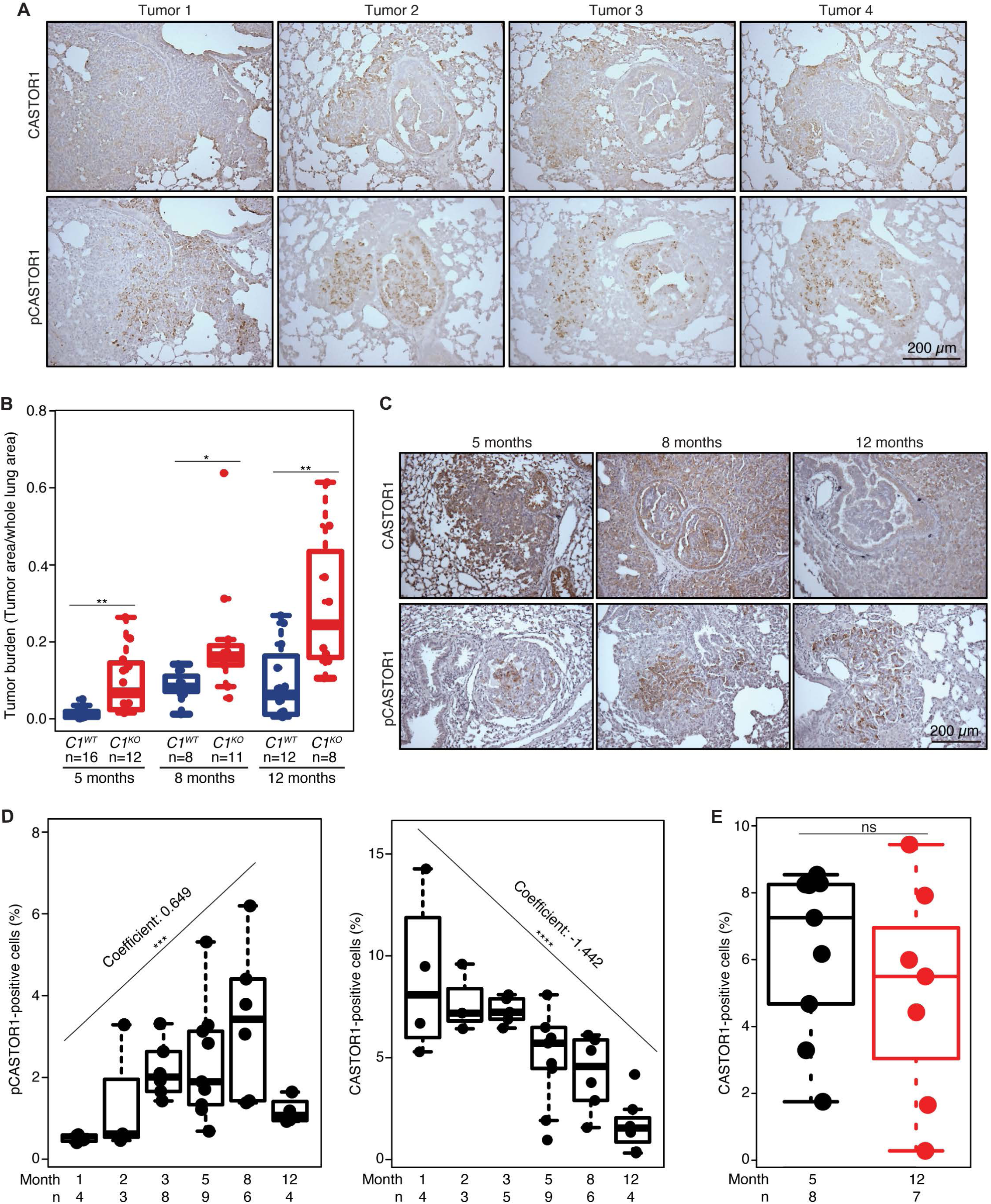
CASTOR1 is inactivated during tumor progression. (A) Inverse correlation of CASTOR1 and pCASTOR1 in *KRAS^G12D^*-induced tumors. Representative IHC staining images of CASTOR1 and pCASTOR1 in consecutively cut sections of lung tumors from four independent *KRAS^G12D^;C1^WT^* mice. (B) Quantification of tumor burden (tumor area/total lung area) in *KRAS^G12D^;C1^WT^* and *KRAS^G12D^;C1^KO^* mice at the indicated time points. Statistical analysis was performed by Student’s t-test. (C) Representative IHC staining images of CASTOR1 and pCASTOR1 in tumors from *KRAS^G12D^;C1^WT^* mice at the indicated time points. (D) Quantification of CASTOR1 and pCASTOR1 levels in tumors from *KRAS^G12D^;C1^WT^* mice at the indicated timepoints. Linear regression analysis was performed without the 12-month time point for pCASTOR1 and with all the time points for total CASTOR1, respectively. (E) Quantification of CASTOR1 levels in adjacent normal lung areas of *KRAS^G12D^;C1^WT^* mice collected at early (5 months) and late (12 months) timepoints. Statistical analysis was performed by Student’s t-test.

We next examined tumor burden in lung tissues collected at various time points. *KRAS^G12D^;C1^KO^* mice exhibited at least a twofold greater tumor burden compared to *KRAS^G12D^;C1^WT^* mice across all examined time points (**Figure 3B**). Furthermore, CASTOR1 levels showed a strong negative correlation with tumor stage, with advanced-stage tumors displaying near-complete loss of CASTOR1 expression (**Figures 3C and 3D**). In contrast, pCASTOR1 levels positively correlated with tumor stage, peaking before CASTOR1 expression was entirely lost in advanced tumors (**Figures 3C and 3D**). Importantly, no changes in CASTOR1 expression were observed in normal lung tissues over the same period (**Figure 3E**), indicating that the time- and tumor-stage-dependent degradation of CASTOR1 is specific to tumor progression.

These results suggest that CASTOR1 inactivation occurs as tumors progress, contributing to enhanced tumor development. The degradation of CASTOR1, driven by AKT-mediated phosphorylation, represents a critical mechanism for its functional inactivation during lung tumor progression.

### CASTOR1 ablation enhances *KRAS^G12D^*-induced genome instability

To investigate whether *CASTOR1* ablation exacerbates *KRAS^G12D^*-induced tumorigenesis by promoting chromosomal instability, we assessed genome instability in MEFs derived from mice with or without *KRAS^G12D^* expression and *CASTOR1* ablation. Chromosomal abnormalities often result in the loss of whole or partial chromosomes, which can be evaluated by measuring micronuclei formation. As anticipated, *KRAS^G12D^* expression alone induced micronuclei in 9% of MEFs compared to 1% in MEFs from *KRAS^WT^* mice (**Figures 4A-B**). Remarkably, *CASTOR1* ablation alone induced micronuclei in 4.9% of the cells. Furthermore, *CASTOR1* ablation significantly enhanced *KRAS^G12D^*-induced micronuclei formation, increasing the percentage to 15% (**Figures 4A-B**).

**Figure 4.**
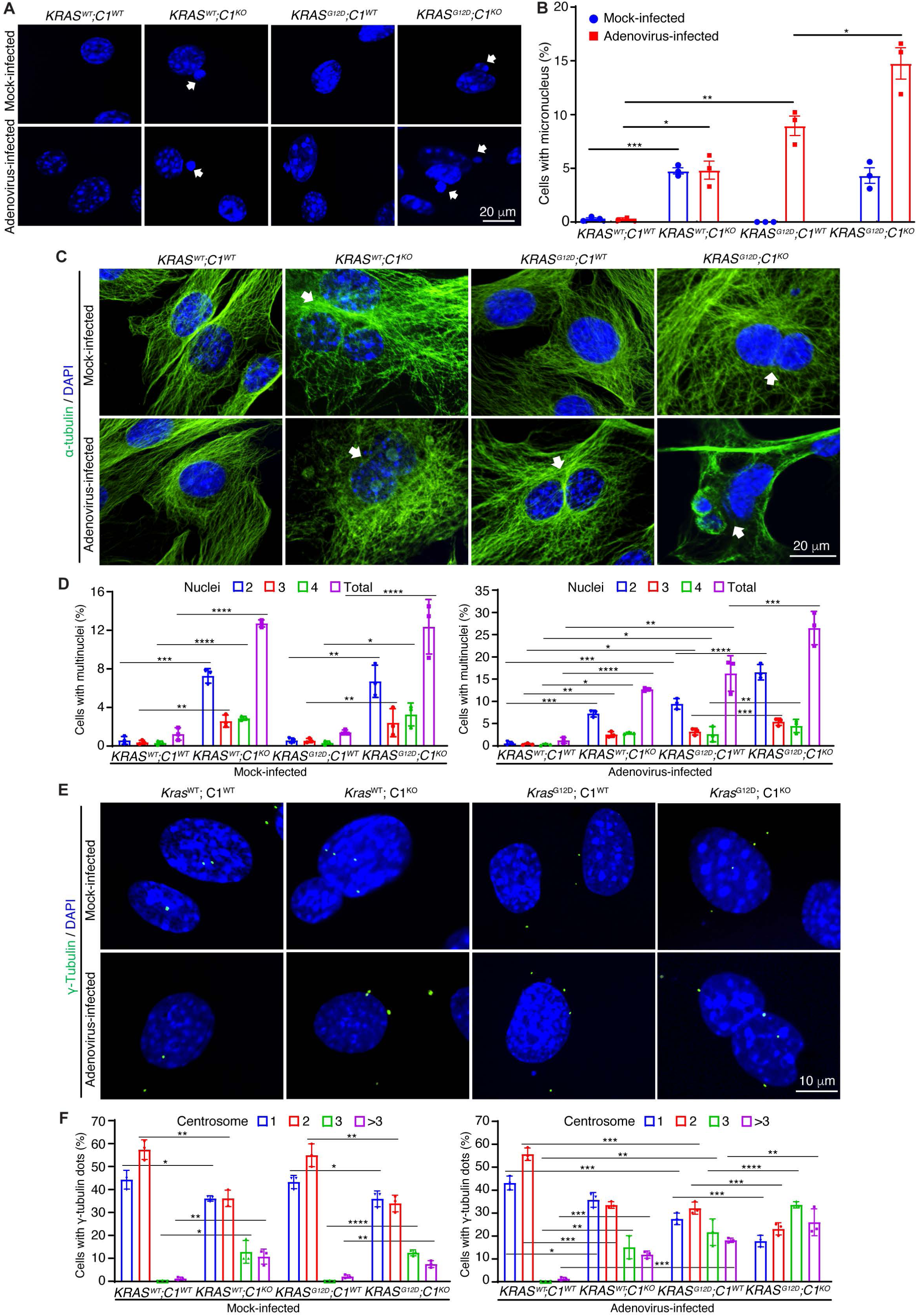
*CASTOR1* ablation enhances *KRAS^G12D^*-induced chromosome instability. (A-B) *CASTOR1* ablation enhances *KRAS^G12D^*-induced micronucleus. Mock- or Adeno-Cre-infected cells were examined for micronucleus at day 3 post-infection by staining for nuclei (DAPI, blue) with representative images shown (A) and quantified (B). Arrow heads indicate representative micronucleus. Statistical analysis was performed by Student’s t-test. (C-D) *CASTOR1* ablation enhances *KRAS^G12D^*-induced multinucleation. Mock- or Adeno-Cre-infected cells were examined for multinucleation at day 3 post-infection by staining for α-tubulin (green) and nuclei (DAPI, blue) with representative images shown (C) and quantified (D). Arrow heads indicate representative micronucleus. Statistical analysis was performed by Student’s t-test. (E-F) *CASTOR1* ablation enhances *KRAS^G12D^*-induced abnormal centrosome duplication. Mock- or Adeno-Cre-infected cells were examined for centrosomes at day 3 post-infection by staining for γ-tubulin (green) and nuclei (DAPI, blue) with representative images shown (E) and quantified (F). Statistical analysis was performed by Student’s t-test.

Chromosomal abnormalities frequently cause aneuploidy and multinucleation due to mitotic dysregulation. Like micronuclei formation, both *CASTOR1* ablation and *KRAS^G12D^* expression independently induced multinucleation, as demonstrated by DAPI and α-tubulin dual staining (**Figures 4C-D**). MEFs with multinucleation often exhibited nuclear atypia, including enlarged, irregularly shaped nuclei and increased cell size. Importantly, *CASTOR1* loss further amplified *KRAS^G12D^*-induced multinucleation.

Centrosome abnormalities are another hallmark of chromosomal instability. Using γ-tubulin staining, we observed that multinucleated cells frequently exhibited aberrant centrosome numbers (>3 centrosomes). *CASTOR1* ablation or *KRAS^G12D^* expression independently increased the proportion of cells with >3 centrosomes from <1% in *KRAS^WT^* MEFs to 28% and 40%, respectively. Strikingly, the combination of *CASTOR1* ablation and *KRAS^G12D^* expression increased this proportion to 57% in MEFs from *KRAS^G12D^;C1^KO^* mice (**Figures 4E-F**).

Together, these findings demonstrate that *CASTOR1* loss is sufficient to induce chromosomal instability and significantly enhances *KRAS^G12D^*-induced genomic instability. This synergistic effect likely contributes to the increased tumorigenic potential of *KRAS^G12D^* upon *CASTOR1* ablation.

### Lung tumors and tumor organoids have heterogeneous responses to KRAS^G12D^ inhibitor MRTX1133, overcome by combination therapy

Recent studies have highlighted resistance to KRAS-specific inhibitors in human lung tumors, limiting their therapeutic efficacy.^25,26^ To investigate this, we examined the responses of *KRAS^G12D^;C1^WT^* and *KRAS^G12D^;C1^KO^* tumors to KRAS^G12D^-specific inhibitor MRTX1133 (**Figure 5A-C**). Among the *KRAS^G12D^;C1^WT^* mice treated with MRTX1133, all tumors responded except for a single nodule exceeding 1×10^5^ μm^2^ (**Figure 5B**). In contrast, *KRAS^G12D^;C1^KO^* tumors exhibited marked resistance, with all five mice developing multiple tumor nodules exceeding 1×10^5^ μm^2^, and three mice exhibiting nodules surpassing 2×10^5^ μm^2^. These results suggest that *CASTOR1* loss confers resistance to KRAS-specific inhibition, exacerbating tumor progression.

**Figure 5.**
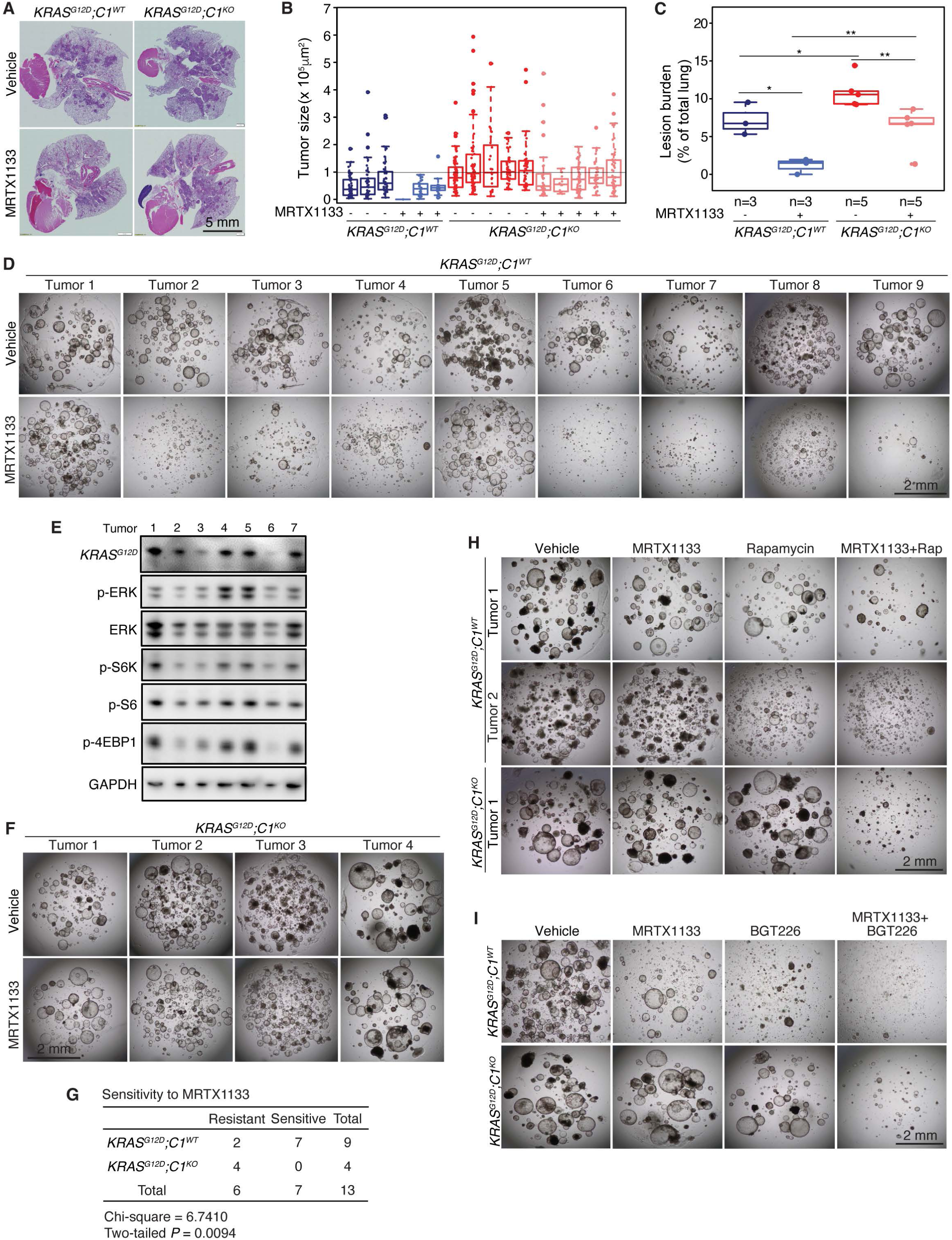
*CASTOR1* ablation enhances resistance to *KRAS^G12D^*-specific inhibitor in tumors and tumor organoids by activating multiple oncogenic pathways. (A-C) *CASTOR1* ablation enhances resistance to *KRAS^G12D^*-specific inhibitor MRTX1133 in tumors. Mice were intranasally inoculated with Adeno-Cre for 4 weeks, then treated with MRTX1133 of vehicle for 4 weeks. Representative H&E images of lungs collected from *KRAS^G12D^;C1^WT^* or *KRAS^G12D^;C1^KO^*mice (A). Tumor size (B) and burden (tumor area/total lung area ×100) (C) were quantified. Data presented as mean +/- standard error of the mean (SEM). Statistical analysis was performed by nested-Anova and Student’s t-test. (D) *KRAS^G12D^;C1^WT^* tumor organoids had heterogeneous response to MRTX1133. Representative pictures of nine *KRAS^G12D^;C1^WT^* organoids after one week of treatment with MRTX1133 or vehicle. (E) Immunoblotting analysis of KRAS^G12D^ protein, mTORC1 downstream targets S6K and 4EBP1, and KRAS downstream target ERK in *KRAS^G12D^;C1^WT^* tumor organoids. (F) *KRAS^G12D^;C1^KO^* tumor organoids were resistant to MRTX1133. Representative pictures of four *KRAS^G12D^;C1^KO^* organoids after one week of treatment with MRTX1133 or vehicle. (G) Comparison of sensitivity to MRTX1133 between organoids derived from *KRAS^G12D^;C1^WT^* and *KRAS^G12D^;C1^KO^* tumors. (H-I) mTORC1 inhibitor Rapamycin and PI3K inhibitor BGT226 sensitized *KRAS^G12D^;C1^WT^* and *KRAS^G12D^;C1^KO^* organoids to MRTX1133. Representative pictures of organoids treated with mono- or combined therapy of MRTX1133 and Rapamycin (H) or BGT226 (I).

To further explore tumor resistance mechanism, we established tumor organoids from both *KRAS^G12D^;C1^WT^* and *KRAS^G12D^;C1^KO^* tumors (**Figure S4A**). Organoids from *KRAS^G12D^;C1^KO^* tumors grew faster than those from *KRAS^G12D^;C1^WT^* tumors, consistent with in vivo results. Analysis of CASTOR1 and pCASTOR1 levels showed a general inverse correlation, mirroring the findings in tumors (**Figure S4B**). We further confirmed activation of the ERK and mTORC1 pathways in these organoids (**Figure S4C-D**). Hence, these organoids recapitulate tumor behavior in mice.

We then tested organoid responses to MRTX1133. The nine independent organoids derived from *KRAS^G12D^;C1^WT^* tumors displayed heterogeneous sensitivity to MRTX1133 with two (Tumors 6 and 7) responding completely, five (Tumors 2, 3, 4, 8 and 9) showing partial responses, and two (Tumors 1 and 5) were completely resistant (**Figure 5D**). Examination of mTORC1 and ERK pathways in in Tumors 1-7 revealed heterogeneous activation, with p-ERK levels correlating with MRTX1133 resistance (**Figure 5E**). In contrast, organoids derived from all four *KRAS^G12D^;C1^KO^* tumors were uniformly resistant to MRTX1133 (**Figure 5F**). Thus, organoids from *KRAS^G12D^;C1^KO^* tumors were significantly more resistant to MRTX1133 than those from *KRAS^G12D^;C1^WT^* tumors (**Figure 5G**). These results further confirm the in vivo results that CASTOR1 loss drives resistance.

Given that mTORC1 activation is commonly observed in human *KRAS*-driven tumors, it may represent a key resistance mechanism to KRAS inhibitors. Dual activation of KRAS and mTORC1 likely enables tumor cells to bypass single-agent therapy. To address this, we tested combination regimens pairing MRTX1133 with either the mTOR inhibitor Rapamycin or the PI3K/AKT dual inhibitor BGT226. Co-targeting KRAS^G12D^ and the PI3K/AKT/mTOR pathways significantly improved therapeutic efficacy in MRTX1133-resistant organoids from *KRAS^G12D^;C1^WT^* tumors (**Figure 5H-I**). Notably, the combination of Rapamycin or BGT226 and MRTX1133 produced a highly synergistic effect in *KRAS^G12D^;C1^KO^* tumor organoids (**Figure 5H-I**), indicating that CASTOR1-deficient tumors rely on KRAS, mTORC1/PI3K/AKT signaling for survival.

These findings highlight mTORC1 hyperactivation as a critical mechanism of resistance to KRAS^G12D^ inhibitors like MRTX1133. They also underscore the therapeutic potential of combination strategies targeting both KRAS and mTORC1 pathways to overcome resistance and improve treatment outcomes.

### CASTOR1 inhibits human lung cancer cell growth by suppressing mTORC1 and other oncogenic pathways

Our findings confirm that CASTOR1 exerts suppressive effects on mTORC1. Furthermore, its ablation leads to the activation and enhancement of *KRAS*-driven oncogenic pathways, including the ERK and PI3K/AKT pathways. This suggests that CASTOR1 may suppress tumorigenesis through additional mechanisms beyond mTORC1 inhibition.

To explore CASTOR1’s alternative functions, we performed *CASTOR1* knockdown using siRNAs in human lung cancer cell lines H2935 and H4006, both of which express high levels of CASTOR1 (**Figure 6A**). Similar to the observations in *KRAS^G12D^*-driven tumors, *CASTOR1* knockdown promoted the proliferation of these lung cancer cells (**Figure 6A**). Immunoblotting analysis revealed increased activation of mTORC1 downstream targets (p-S6K and p-4EBP1) as well as the AKT and ERK pathways. Conversely, overexpression of *CASTOR1* in lung cancer cell lines H647, H1648, and H1975, which express low levels of CASTOR1, significantly inhibited their growth (**Figure 6B**). Immunoblotting further confirmed suppression of mTORC1 activity and downregulation of AKT and ERK pathways following CASTOR1 overexpression (**Figure 6B**).

**Figure 6.**
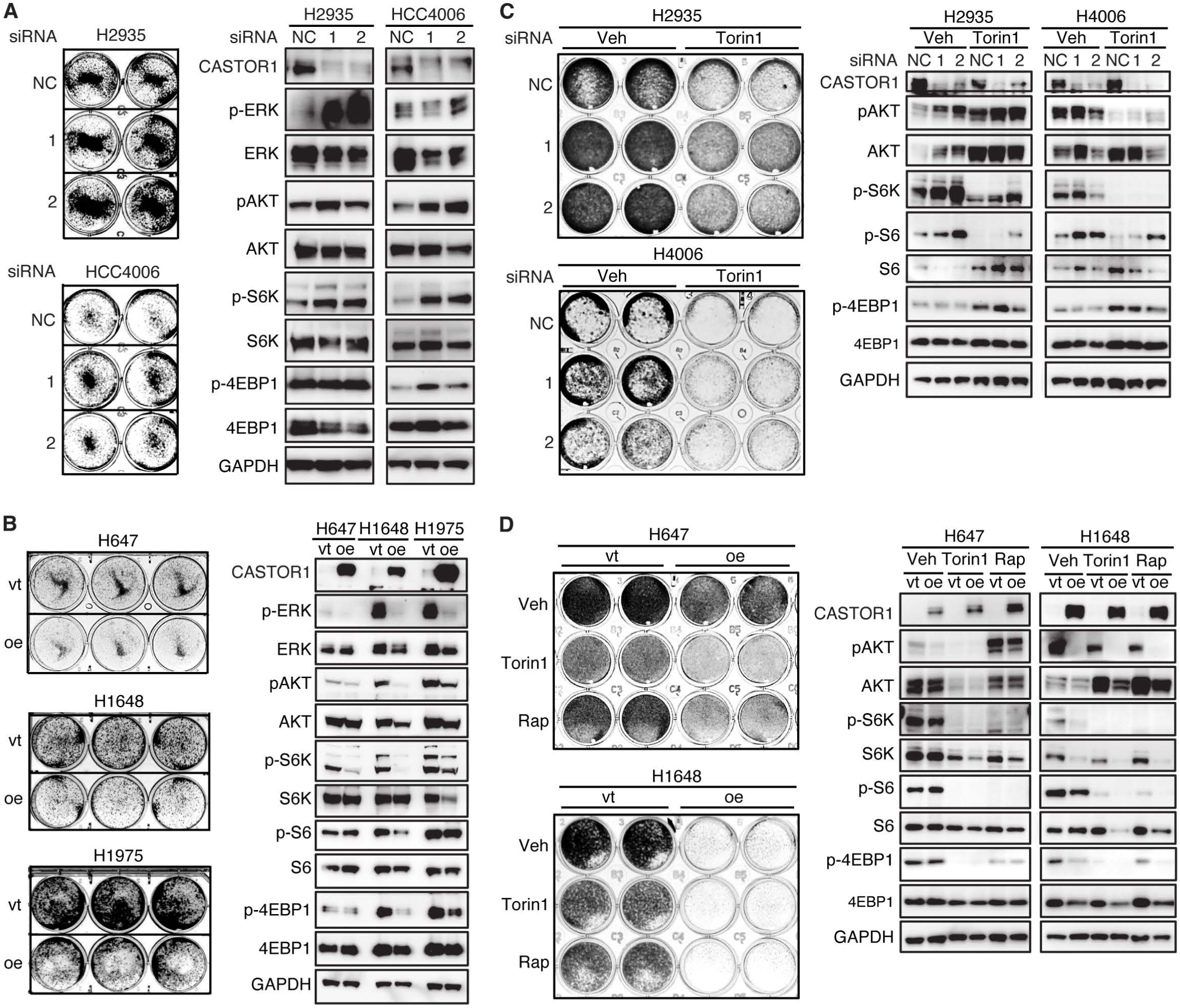
CASTOR1 mediates proliferation and resistance to mTORC1 inhibitors. (A) Knockdown of *CASTOR1* enhanced cell proliferation of human lung cancer cell lines. Clonogenic assay and immunoblotting of human LUAD cell lines following siRNA knockdown of *CASTOR1*. (B) Overexpression of *CASTOR1* inhibited cell proliferation of human lung cancer cell lines. Clonogenic assay and immunoblotting of human LUAD cell lines following overexpression of *CASTOR1*. (C) Knockdown of *CASTOR1* enhanced resistance to mTORC1 inhibitor Torin1 in human lung cancer cell lines. Clonogenic assay and immunoblotting of human LUAD cell lines treated with Torin1 following siRNA knockdown of *CASTOR1*. (D) Overexpression of *CASTOR1* sensitized human lung cancer cell lines to mTORC1 inhibitors Torin1 and Rapamycin. Clonogenic assay and immunoblotting of human LUAD cell lines with mTOR inhibitors Torin1 and Rapamycin following overexpression of *CASTOR1*.

### CASTOR1 enhances lung cancer cell sensitivity to mTORC1 inhibitors

Given CASTOR1’s regulatory effects on mTORC1 and other oncogenic pathways, we investigated whether CASTOR1 modulation affects lung cancer cell responses to mTORC1 inhibitors. In H2935 and H4006 cells, *CASTOR1* knockdown not only conferred a growth advantage but also partially rescued the cells from the cytotoxic effect of mTORC1 inhibitor Torin1 (**Figure 6C**). While immunoblotting analysis revealed increased activation of mTORC1 downstream targets (p-S6, p-S6K and p-4EBP1) as well as the AKT following *CASTOR1* knockdown, treatment with Torin1 enhanced AKT activation, which was likely due to the negative feedback of the mTORC1 inhibition (**Figure 6C**). Conversely, *CASTOR1* overexpression in H647 and H1648 cells significantly suppressed their proliferation and enhanced sensitivity to Torin1 and Rapamycin (**Figure 6D**). *CASTOR1* overexpression suppressed mTORC1 activity and this effect synergized with the mTORC1 inhibitors (**Figure 6D**).

These findings suggest that the combination of CASTOR1 activation and mTORC1 inhibition provides an additive effect, potentially suppressing both the mTORC1 pathway and other oncogenic pathways. This highlights the therapeutic potential of dual targeting strategies for treating lung cancer.

## Discussion

The PI3K/AKT/mTOR pathway is well-known for its critical role in tumorigenesis and disease progression in NSCLC.^27^ Somatic mutations and amplifications of *PIK3CA* are frequently found in NSCLC patients, and the AKT pathway is activated in a substantial proportion of cases.^28,29^ The mTOR pathway is activated in a significant proportion of NSCLC tumors, with elevated p-mTOR levels detected in up to 90% of adenocarcinomas, 60% of large cell carcinomas, and 40% of squamous cell carcinomas.^27,30,31^ Furthermore, activation of downstream targets of mTORC1, such as S6K and 4EBP1, is also common and correlates with poor prognosis in early-stage NSCLC.^15,32,33^ These findings underscore the clinical relevance of the mTORC1 pathway and its components as critical regulators of lung cancer progression.

In this study, we identify CASTOR1 as a novel tumor suppressor that functions through the modulation of multiple oncogenic pathways, including but not limited to the mTORC1 signaling cascade. As an arginine sensor, CASTOR1 plays a pivotal role in regulating mTORC1 activity by modulating its lysosomal translocation.^17,34^ Beyond this canonical role, our results reveal that CASTOR1 also inhibits *KRAS*-induced oncogenic pathways, positioning it as a central regulator of tumor progression. This dual role of CASTOR1 highlights its importance in the tumor development, where its loss can exacerbate the effects of oncogenic *KRAS* mutations.

### *CASTOR1* as a novel concept in tumor suppressor biology

Our findings introduce a novel concept in tumor suppressor biology. Unlike traditional tumor suppressors, which predispose cells to tumor initiation when inactivated, *CASTOR1* deletion does not lead to spontaneous tumor formation in mice. Examination of TCGA database and other database did not identify association of *CASTOR1* amplification, deletion or mutation with any types of cancer. Instead, its loss appears to accelerate tumor progression in the presence of additional oncogenic insults, such as mutant *KRAS*. This was evident in our double transgenic *KRAS^G12D^;C1^KO^* mouse model, where the absence of CASTOR1 synergized with mutant *KRAS^G12D^* to drive lung cancer development at a significantly accelerated rate compared to *KRAS* activation alone in *KRAS^G12D^;C1^WT^* mice.

Significantly, CASTOR1 is inactivated during the development of *KRAS-*driven lung cancer. Notably, CASTOR1 inactivation occurs via AKT-mediated phosphorylation at serine 14, leading to its proteasomal degradation^18^. This represents a novel mode of tumor suppressor regulation that is distinct from the genetic mutations or deletions typically seen in other tumor suppressors. Immunohistochemical analysis revealed an inverse correlation between CASTOR1 expression and tumor progression stages, suggesting that CASTOR1 inactivation plays a critical role during tumor development. These results are in agreement with the observation that the reduced expression of this protein is strongly correlated with poor prognosis in many cancer types as well^18^ and that pCASTOR1 levels predict poor prognosis in early-stage male patients with *KRAS* mutations.^19^ These findings highlight the unique characteristics of CASTOR1 as a tumor suppressor and its potential as a therapeutic target.

### Mechanisms of CASTOR1 tumor suppression

Our results show that CASTOR1 likely suppresses tumorigenesis through inhibiting multiple oncogenic pathways, including but not limited to the mTORC1 signaling pathway. It has been documented that the crosstalk and compensation between mTORC1 and KRAS pathway is very sophisticated.^35^ Our findings demonstrate that *CASTOR1* ablation alone is not only sufficient to activate mTORC1, AKT and ERK pathways but also enhances KRAS-activation of these pathways. While the specific mechanisms of these effects are unclear, they are likely important in lung cancer as *CASTOR1* is inactivated during lung cancer development.^18,19^ Furthermore, *KRAS* mutations underlie 30% of NSCLC and predict negative outcomes in early- and late-stage disease.^36,37^

Abnormal activation of oncogenic pathways often leads to chromosome instability, a key factor contributing to tumor progression and aggressive cancer phenotypes in vivo.^38^ Hence, genome instability, characterized by chromosomal aberrations, aneuploidy, and DNA damage accumulation, has long been recognized as a hallmark of cancer, driving tumor evolution and resistance to therapy. Lung cancer has high genomic instability,^39^ which can accelerate the acquisition of genetic diversity and promote the formation of various cancer characteristics. Our findings demonstrate that *CASTOR1* ablation induces and enhances *KRAS*-driven genome instability. By synergizing with KRAS-induced oncogenic pathways, *CASTOR1* loss amplifies genomic instability, most likely by exacerbating KRAS-mediated genomic instability through its downstream effects, thereby accelerating tumorigenesis. This suggests that CASTOR1 functions as a gatekeeper of genomic integrity, preventing the accumulation of deleterious mutations during tumor development. Unlike classical tumor suppressors, whose inactivation predisposes cells to transformation, CASTOR1 appears to be selectively downregulated during tumor progression rather than at the initiation stage. This novel concept of tumor suppressor inactivation during disease progression highlights a previously underexplored paradigm in cancer biology.

### Mechanisms of resistance in human lung tumors

The dysregulation of multiple oncogenic pathways, including the mTORC1 pathway, contributes to therapeutic resistance, particularly to KRAS-specific inhibitors. Our study and others have shown that mTORC1 activation, as indicated by elevated levels of p-S6K and p-4EBP1, is a common mechanism of resistance to KRAS inhibitors.^23,26^ The interplay between mTORC1 and KRAS signaling is particularly significant in NSCLC, where compensatory activation of mTORC1 often diminishes the efficacy of KRAS-targeted therapies.

Using lung cancer organoids derived from *KRAS^G12D^*-driven tumors, we further demonstrated heterogeneity in the response to MRTX1133, revealing the intrinsic resistance in these tumors. While 2 of 9 organoids were sensitive to the drug, the remaining 7 exhibited varying degrees of resistance, which correlated with higher mTORC1 activity. In vivo results show that *KRAS^G12D^;C1^KO^* tumors, which have hyperactivation of the mTORC1 pathway, are more resistant to MRTX1133 than *K^G12D^;C1^WT^* tumors. This suggests that elevated mTORC1 signaling serves as a resistance mechanism, reinforcing the need for combination therapies targeting both KRAS and mTORC1 pathways. An extra layer of complexity arises from the cross-regulation between the two pathways is the observation of activated ERK as it interacts with multiple nodes of the PI3K/AKT/mTOR pathway, including its phosphorylation of TSC2.^40,41^ Our findings provide compelling evidence that combination therapy with KRAS-specific inhibitors and mTORC1 inhibitors, and likely inhibitors targeting PI3K/AKT and ERK pathways can effectively overcome resistance in lung cancer. In organoids, the addition of Rapamycin, a mTORC1 inhibitor, and BGT226, a PI3K inhibitor, to MRTX1133 restored sensitivity in resistant organoids. These results align with previous reports highlighting 4EBP1 is a convergent target of PI3K/AKT/mTOR and RAF/MEK/ERK pathways that integrates their functions at the level of regulation of cap-dependent translation.^42^ Indeed, inhibition of either PI3K/mTOR or ERK signaling often relieves feedback inhibition of the other and causes its activation.^43,44^ Given that previous strategies combining MEK and AKT inhibition proved to be poorly tolerated in *KRAS^G12D^* mutant NSCLC patients and not appropriate for treatment of these patients,^45,46^ combination therapy targeting both pathways not only mitigates resistance but also enhances therapeutic efficacy, underscoring its potential as a clinical strategy.

### CASTOR1 activation as a therapeutic approach

In addition to combination therapy, activation of CASTOR1 represents a promising therapeutic strategy. As a natural inhibitor of mTORC1 and other oncogenic pathways, restoring CASTOR1 function could enhance the response to KRAS-specific therapies. AKT-mediated degradation of CASTOR1 represents a key regulatory mechanism that could be targeted to stabilize its expression. For example, pharmacological inhibitors of AKT or gene-editing strategies designed to block CASTOR1 phosphorylation at serine 14 and RNF167-mediated ubiquitination could restore its tumor-suppressive function. Restoring or mimicking CASTOR1 activity not only offers the potential to suppress mTORC1 and KRAS-driven oncogenic pathways but may also delay or prevent the emergence of resistance to KRAS-specific inhibitors. This dual benefit positions CASTOR1 as a promising target for therapeutic intervention in lung cancers driven by *KRAS* mutations and mTORC1 activation.

## Conclusion

This study establishes *CASTOR1* as a novel tumor suppressor gene with unique regulatory mechanisms and highlights its critical role in modulating both mTORC1 and KRAS oncogenic pathways. The discovery of CASTOR1’s degradation during tumor progression introduces a new paradigm in tumor suppressor biology and underscores its potential as a therapeutic target. Furthermore, our findings provide strong preclinical evidence supporting combination therapy with KRAS-specific inhibitors and mTORC1 inhibitors, as well as strategies to restore CASTOR1 activity, to improve outcomes in NSCLC patients with *KRAS* mutations.

## Materials and Methods

### *CASTOR1* knockout mice and *KRAS^G12D^* transgenic mice

*CASTOR1* KO mice (*CASTOR1^KO^* or *C1^KO^*) were generated by artificial insemination conducted by The Mouse Embryo Services (MES) Core in the Department of Immunology at the University of Pittsburgh. In brief, an aliquot of cryopreserved sperm of C57BL/6N-*CASTOR1tm1.1^(KOMP)Vlcg^*/MbpMmucd (Stock# 046916-UCD) was purchased from the Mouse Biology Program (MBP) at the University of California Davis and delivered to MES Core for artificial insemination. All newborn pups were genotyped by standard genotyping PCR following the protocol provided by MBP using the Platinum™ SuperFi™ PCR Master Mix (Thermo, cat. No. 12358050). The primers used include CASTOR1-Reg-Lac-F (5’ ACTTGCTTTAAAAAACCTCCCACAC 3’) and CASTOR1-Reg-R1 (5’ TAAACTGCTTGAAAAGCCAAGGACGG 3’) for *CASTOR1^KO^* mice; and CASTOR1-WT-F (5*’ AAGTTCT TCAGCCTGACTGAGACTCC* 3’) and CASTOR1-WT-R (5’ GCCTGGGACTCATATTCA GCTCC 3’) for *CASTOR1^WT^* mice. The absence of *CASTOR1* expression in *CASTOR1^KO^* mice was validated by both RT-qPCR and immunoblotting (**Figure S1C-D**). *KRAS^LSL-G12D^* or *KRAS^G12D^* mouse was originally generated by the Jackson Laboratory (008179). All newborn pups were genotyped by protocol provided by the vendor with the following primers, including forward primers KRAS-WT-F (5’ TGTCTTTC CCCAGCACAGT 3’) and KRAS-Mut-F (5’ GCAGGTCGAGGGACCTAATA 3’) for KRAS and *KRAS^G12D^* mice, respectively, and reverse primer KRAS-Common-R (5’ CTGCATAGTACGCTATACCCTGT 3’) for both mice. *KRAS^G12D^*;*C1^KO^* were established by crossing *KRAS^G12D^* and *CASTOR1^KO^* mice.

Cre-expressing adenovirus (Adeno-Cre) was purchased from the Viral Vector Core Facility (University of Iowa, Iowa City, IA). The expression of *KRAS^G12D^* was induced with intranasal administration of Adeno-Cre (2×10^7^ IFU/mouse).^47^ Mice were maintained in pathogen-free conditions with free access to food and water. Mice that developed a tumor-unrelated illness during the experiment were euthanized and excluded from the study. Lung as well as other organs were collected at various time points post-injection of adenovirus. All tissues were fixed in 10% formalin overnight, transferred to 70% ethanol, and subsequently embedded in paraffin.

For in vivo drug treatment, *KRAS^G12D^*;*C1^WT^* and *KRAS^G12D^*;*C1^KO^* mice were exposed to Adeno-Cre through intranasal injection as described above. At week 4 after injection, mice were randomized into either vehicle or MRTX1133 treatment groups. MRTX1133 was freshly formulated and administered at 3 mg/kg by intraperitoneal injection twice daily and the treatment continued for 4 weeks. All mice were then euthanized and the lung tissues were collected to assess the drug effect.

All animal experiments were performed in accordance with National Institute of Health guidelines and were approved by the University of Pittsburgh Animal Care and Use Committee (Protocol #: 21079422 and 24075063).

### Mouse embryonic fibroblasts and induction of *KRAS^G12D^* expression

Mouse embryonic fibroblasts (MEFs) were isolated from female mouse. Briefly, MEFs were derived from mouse embryos at gestational day E11.5–E13.5 via uterine dissection. Each embryo was rinsed with 1× PBS (pH 7.4), and the internal organs were carefully removed. The embryonic bodies were transferred into a clean 15 mL tube containing 3 mL of 0.25% trypsin-EDTA and homogenized by pipetting. The resulting tissue homogenate was incubated at 37°C for 30 minutes, then further dissociated by pipetting. The suspension was divided evenly into 10 cm tissue culture dishes and cultured in Dulbecco’s Modified Eagle Medium (DMEM) supplemented with 20% fetal bovine serum (FBS, Invitrogen) and 1% penicillin-streptomycin. Early-passage MEFs (passages 1-10) were used for all experiments, and cells derived from at least three different embryos were analyzed.

The expression of *KRAS^G12D^* in MEFs was induced by Adeno-Cre infection. MEFs from each embryo were split into two groups: one infected with Adeno-Cre to induce *KRAS^G12D^* expression and the other mock-infected as a control. Confluent MEFs cultured in 6-well plates were infected with Adeno-Cre at a multiplicity of infection (MOI) of 50. After a 2-hour adsorption period, unabsorbed virus particles were removed by washing the cells twice with DMEM. Cells were then maintained in fresh DMEM containing 20% FBS. At day 3 post-infection, cells were either cryopreserved or harvested for *KRAS^G12D^* expression analysis via immunoblotting and immunofluorescence staining.

### Generation of tumor organoids

Tumor organoids were generated from tumors of *KRAS^G12D^*;*C1^WT^* and *KRAS^G12D^*;*C1^KO^* mice as previously described.^48,49^ Briefly, tumors were excised and sliced into small fragments. Tumor fragments were assessed for tumor cell content before organoid generation. The fragments were washed with cold PBS, and the supernatant was collected by allowing the fragments to settle under normal gravity for 1 minute. Single cells from the supernatant were then pelleted by centrifugation, washed with PBS, and resuspended in growth factor-reduced, phenol red-free Matrigel (Corning #356231) on ice. Cells were plated in 24-well plates at a density of 15,000 cells per 50 µL Matrigel. The plates were incubated at 37°C to allow Matrigel polymerization. After 15 minutes, 500 µL of organoid complete medium (OCM) supplemented with 50 ng/mL murine EGF was added to each well. The medium was refreshed every 2-3 days, and organoid growth was monitored daily using an inverted microscope.

For drug treatment assays, organoids were treated with drugs at indicated dosages as following: MRTX1133 10 nM (ChemGood, cat.no. C1420), Rapamycin 1 μM (MCE, cat.no. HY10219), and BGT226 10 nM (MCE, cat.no. 13334). OCM was changed every other day with freshly prepared drugs added. At the end point, organoids are either lyzed in 1x sample buffer for immunoblotting analysis or fixed in 1% neutral buffered formalin for immunohistochemistry (IHC) analysis.

For passaging, organoids embedded in Matrigel were mechanically disrupted using a P1000 pipette tip, collected, washed with OCM, resuspended, and re-plated at 50 µL Matrigel per well. For cryopreservation, organoids were disrupted with a P1000 pipette fitted with cut-off tips, transferred into a 15 mL Falcon tube, washed with 5 mL OCM, and centrifuged at 200 × g for 2 minutes. The pellet was resuspended in OCM containing 10% dimethyl sulfoxide (DMSO), and 1 mL aliquots were transferred into cryovials. These cryovials were placed in a Corning CoolCell® container (Corning, NY) and stored at −80°C for initial freezing, then transferred to vapor-phase liquid nitrogen for long-term storage.

### Immunofluorescence staining

For immunofluorescence staining, MEFs were plated on glass coverslips and cultured overnight. The cells were fixed and permeabilized with 100% methanol at −20°C for 10 minutes. Blocking was performed with 3% BSA for 60 minutes at room temperature. Primary antibodies, including anti-KRAS (CST, E4K9L, #91054), anti-KRAS^G12D^ (Thermo Fisher, HL10, MA5-36256), anti-α-tubulin (Sigma-Aldrich, T6199), and anti-γ-tubulin (Sigma-Aldrich, T6557) were diluted 1:500 in 3% BSA and incubated with MEFs for 1 hour at room temperature. Following primary antibody incubation, the cells were washed with PBS and incubated with either goat anti-mouse (Thermo Fisher, A11001) or goat anti-rabbit (Thermo Fisher, A11008) secondary antibodies conjugated with FITC, diluted 1:40 in 3% BSA for 45 minutes at room temperature. After secondary antibody incubation, cells were washed in PBS and counterstained with 4,6-diamidino-2-phenylindole (DAPI, Sigma-Aldrich, D9542) for nuclear visualization.

To assess chromosome instability in MEFs, micronuclei were detected by staining the cells with DAPI at a concentration of 1 μg/mL for 30 minutes at room temperature. Immunofluorescence images were captured using a fluorescence microscope (Nikon Microscope, Melville, NY, USA).

### Immunohistochemistry

Tissues were fixed in 10% neutral buffered formalin for up to 24 hours and transferred to 70% ethanol for storage. IHC was performed on formalin-fixed, paraffin-embedded (FFPE) lung sections, which were cut into 5-μm slices for hematoxylin and eosin (H&E), and Ki67 staining as previously described.^50^ Sections were deparaffinized by heating at 95°C for 10 minutes, followed by xylene rinsing, and antigen retrieval was performed in citrate buffer (pH 6.0) using a microwave at full power for 20 minutes. Endogenous peroxidase activity was quenched with 3% hydrogen peroxide, and nonspecific binding was blocked using 5% bovine serum albumin (BSA). Slides were incubated with primary antibodies overnight at 4°C, then secondary antibodies at room temperature for 1 hour, and signals were detected using horseradish peroxidase (HRP)- based diaminobenzidine (DAB) substrate, with hematoxylin used for counterstaining. Primary antibodies include CASTOR1 (Abclonal, cat. no. A2071C, 1:50 dilution), p-ERK (Cell Signaling Technology, cat. no.4370, 1:100 dilution), p-4EBP1 (Cell Signaling Technology, cat. no.2855, 1:100), p-S6K (Abclonal, cat. no.AP0502, 1:100 dilution), p-S6 (Cell Signaling Technology, cat. no.2211, 1:100), and pCASTOR1 (1:100). The pCASTOR1 antibody (1:100 dilution) was previously described.^19^

### Cell culture and clonogenic assay

Human lung cancer cell lines HCC2935, HCC4006, H647, H12648 and H1975 were cultured in RPMI-1640 medium supplemented with 10% FBS. For transfection, lipofectamine 2000 (Thermo Fisher Scientific, cat.no.11668030) was used for transient transfection of plasmids, and RNAimax (Thermo Fisher Scientific, cat.no.13778100) was used for small interfering RNA tansfection following the manufacturer’s instructions. For drug treatment, medium containg freshed fomulated Rapamycin (1 μM, MCE, cat.no. HY10219) or Torin1 (0.2 μM, MCE, cat.no. HY13003) was changed every 3 days. At designated time points, the medium was removed, and the cells were fixed using a fixation solution composed of acetic acid and methanol (1:7) for 8 minutes. The fixation solution was replaced with 0.5% crystal violet, and the cells were incubated at room temperature for 2 hours. Excess crystal violet was carefully rinsed off under tap water, and the dishes were air-dried overnight at room temperature before analysis.

### RNA isolation and quantitative reverse transcription real-time PCR (RT-qPCR)

Total RNA was isolated using Trizol reagent following the manufacturer’s instructions (Invitrogen). Quantitative reverse transcrption real-time PCR (RT-qPCR) was performed to measure relative mRNA levels of target genes using SYBR® Green (Thermo Fisher Scientific) on a CFX Real-Time PCR System (Bio-Rad). Sample loading was normalized with *β-actin* as the internal control. PCR reactions were carried out under the following conditions: an initial denaturation step at 95°C for 10 minutes, followed by 40 cycles of 95°C for 15 seconds and 60°C for 45 seconds. Specificity of the amplification products was verified through dissociation curve analysis. Transcript levels were quantified using the comparative Ct method, and the fold-change of target gene expression normalized to the internal control was calculated using the formula 2-△△CT. All data represent the mean of three independent replicates. The primers used include: *Mus musculus CASTOR1* (Forward: 5’ TGCGAGTACTGAGCATTGCC 3’; Reverse: 5’ GCTGAAGAACTTGCACCGACTG 3’); *Mus musculus β-Actin* (Forward: 5*’ CCCTGAAGTACCCCATTGAA* 3’; Reverse: 5’ *GGGGTGTTGAAGGTCTCAAA* 3’); *Homo sapiens CASTOR1* (Forward: 5’ -GCTCCATCACGTTCTTTGCC-3’; Reverse: 5’ CCACATTCATCAAAGCCCAGG-3’); and *Homo sapiens GAPDH* (Forward: 5’ AATCCCATCACCATCTTC 3’; Reverse: 5’ AGGCTGTTGTCATACTTC 3’).

### Preparation of total lysates and immunoblotting analysis

Total protein lysates were prepared using RIPA buffer supplemented with a protease inhibitor cocktail (Thermo Scientific, cat. no.186182). Proteins were resolved by sodium dodecyl sulfate-polyacrylamide gel electrophoresis (SDS-PAGE) and subsequently transferred onto nitrocellulose membranes (Millipore, cat. no. GE10600004). Membranes were incubated with primary antibodies overnight at 4°C. The next day, membranes were washed three times with PBS and incubated with goat anti-rabbit HRP conjugated IgG (Cell Signaling Technology, cat. No. 7074S) or horse anti-mouse IgG HRP conjugated IgG (Cell Signaling Technology, cat. No.7076) for 1 hour at room temperature. Protein signals were detected using an SuperSignal^TM^ West Femto Maximum Sensitivity Substrate (Thermo Scientific, cat. no. 34094) and visualized with the Chemidoc MP Imaging System (Bio-Rad). The primary antibodies used in the analysis included: anti-CASTOR1 (Abclonal, cat. no. A2071C), anti-AKT (Cell Signaling Technology, cat. no. 4691), anti-p-AKT-S473 (Cell Signaling Technology, cat. no. 4060), anti-pAKT-T308 (Cell Signaling Technology, cat. no. 4056), anti-S6K (Cell Signaling Technology, cat. no. 97596), anti-4EBP1 (Cell Signaling Technology, cat. no. 9644), anti-p-4EBP1 (Cell Signaling Technology, cat. no. 2855), anti-S6 (Cell Signaling Technology, cat. no. 2217), anti-p-S6 (Cell Signaling Technology, cat. no. 2211), anti-p-ERK (Cell Signaling Technology, cat. no. 4370), anti-RAS (Cell Signaling Technology, cat. no. 91054), anti-*KRAS^G12D^* (Cell Signaling Technology, cat. no. 14429), and anti-GAPDH (Cell Signaling Technology, cat. no. 5174).

### Statistical analysis

All statistical analyses were conducted using GraphPad Prism 6.0 (GraphPad Software, Inc.). Experimental results are presented as the mean ± standard deviation (SD). For comparisons involving multiple groups, data were analyzed using one-way ANOVA followed by Tukey’s post hoc test. A p-value of less than 0.05 (p < 0.05) was considered statistically significant. All data were derived from at least three independent experiments to ensure reproducibility and reliability.

## Resource Availability

### Lead contact

Further information and requests for resources and reagents should be directed to and will be fulfilled by the lead contact, Shou-Jiang Gao.

### Materials availability

Proprietary material is available upon request from the authors.

## Acknowledgments

We thank members of Gao laboratory for assistance and helpful comments. Funding source: This work was supported by grants from National Institutes of Health (CA096512, CA278812, CA284554, CA124332 and CA291244 to S.-J. Gao), and in part by award P30CA047904.

## Author Contributions

Conceptualization, S.-J.G.; methodology, X.W. and L.D.; investigation, X.W., L.D., S.Y.S., S.K.L., L.P.C., T.T.L., M.T.W., A.P., Y.F.H. and S.-J.G.; writing – original draft, X.W., L.D. and S.-J.G.; writing – review & editing, X.W., L.D., S.Y.S., S.K.L., L.P.C., T.T.L., M.T.W., A.P., Y.F.H. and S.-J.G.; funding acquisition, S.-J.G.; supervision, S.-J.G.

## Supplemental Information

Supplemental information can be found online at:.

## Supplemental Materials

**Supplementary figure 1. Full-body knockout of *CASTOR1* in C57/BL6 mouse.**

(A) Schematic illustration of the generation of full-body CASTOR1-null mouse.

(B) PCR genotyping of *CASTOR1*-null mouse. WT: wild-type; Het: heterozygous; KO: knockout.

(C and D) Examination of *CASTOR1* expression by RT-qPCR (C) and immunoblotting

(D) in WT and KO mice.

**Supplementary figure 2. Generation of novel *KRAS^G12D^* mice with *CASTOR1* deletion.**

(A) Schematic illustration of crossing strategy between *KRAS^WT^;C1^KO^* and *KRAS^G12D^;C1^WT^* lines and the offspring with *KRAS^WT^;C1^WT^*, *KRAS^WT^;C1^KO^*, *KRAS^G12D^;C1^WT^* and KRAS^G12D^;C1^KO^ genotypes.

(B) PCR genotyping of *KRAS^WT^;C1^WT^*, *KRAS^WT^;C1^KO^*, *KRAS^G12D^;C1^WT^* and *KRAS^G12D^;C1^KO^* mice.

**Supplementary figure 3. Detection of KRAS and KRAS^G12D^ proteins by immunofluorescence analysis.**

Passage 2 MEFs isolated from *KRAS^WT^;C1^WT^*, *KRAS^WT^;C1^KO^*, *KRAS^G12D^;C1^WT^* and *KRAS^G12D^;C1^KO^* mice were examined for the expression of KRAS (green) and KRAS^G12D^ (green speckles) proteins following mock or Adeno-Cre infection for 3 days. Nuclei were identified by DAPI staining (blue).

**Supplementary figure 4. Generation of *KRAS^G12D^;C1^WT^* and *KRAS^G12D^;C1^KO^* tumor organoids that recapitulate tumor behavior in mice**

(A) Representative pictures of *KRAS^G12D^;C1^WT^* and *KRAS^G12D^;C1^KO^* tumor organoids at day 1, 3 and 6 post-seeding.

(B) Representative IHC staining images of CASTOR1 and pCASTOR1 in three tumor organoids from *KRAS^G12D^;C1^WT^* mice.

(C) Representative immunoblotting analysis of mTORC1 downstream targets S6K and 4EBP1, KRAS downstream target ERK, and their phosphorylated forms in *KRAS^G12D^;C1^WT^* and *KRAS^G12D^;C1^KO^* tumor organoids.

(D) Representative IHC images of p-S6K and p-ERK in *KRAS^G12D^;C1^WT^* and *KRAS^G12D^;C1^KO^* tumor organoids (left panels) and quantifications (right panels).

## References

1 Bray, F. et al. Global cancer statistics 2022: GLOBOCAN estimates of incidence and mortality worldwide for 36 cancers in 185 countries. CA Cancer J Clin 74, 229–263, doi:10.3322/caac.21834 (2024).

2 Siegel, R. L., Kratzer, T. B., Giaquinto, A. N., Sung, H. & Jemal, A. Cancer statistics, 2025. CA Cancer J Clin 75, 10–45, doi:10.3322/caac.21871 (2025).

3 Campbell, J. D. et al. Distinct patterns of somatic genome alterations in lung adenocarcinomas and squamous cell carcinomas. Nat Genet 48, 607–616, doi:10.1038/ng.3564 (2016).

4 Chen, Z., Fillmore, C. M., Hammerman, P. S., Kim, C. F. & Wong, K. K. Non-small-cell lung cancers: a heterogeneous set of diseases. Nat Rev Cancer 14, 535–546, doi:10.1038/nrc3775 (2014).

5 Cox, A. D., Fesik, S. W., Kimmelman, A. C., Luo, J. & Der, C. J. Drugging the undruggable RAS: Mission possible? Nat Rev Drug Discov 13, 828–851, doi:10.1038/nrd4389 (2014).

6 Ostrem, J. M. & Shokat, K. M. Direct small-molecule inhibitors of KRAS: from structural insights to mechanism-based design. Nat Rev Drug Discov 15, 771–785, doi:10.1038/nrd.2016.139 (2016).

7 Simanshu, D. K., Nissley, D. V. & McCormick, F. RAS Proteins and Their Regulators in Human Disease. Cell 170, 17–33, doi:10.1016/j.cell.2017.06.009 (2017).

8. Cancer Genome Atlas Research, N. Comprehensive molecular profiling of lung adenocarcinoma. Nature 511, 543–550, doi:10.1038/nature13385 (2014).

9 Canon, J. et al. The clinical KRAS(G12C) inhibitor AMG 510 drives anti-tumour immunity. Nature 575, 217–223, doi:10.1038/s41586-019-1694-1 (2019).

10 Hallin, J. et al. The KRAS(G12C) Inhibitor MRTX849 Provides Insight toward Therapeutic Susceptibility of KRAS-Mutant Cancers in Mouse Models and Patients. Cancer Discov 10, 54–71, doi:10.1158/2159-8290.CD-19-1167 (2020).

11 Skoulidis, F. et al. Co-occurring genomic alterations define major subsets of KRAS-mutant lung adenocarcinoma with distinct biology, immune profiles, and therapeutic vulnerabilities. Cancer Discov 5, 860–877, doi:10.1158/2159-8290.CD-14-1236 (2015).

12 Sanaei, M. J., Razi, S., Pourbagheri-Sigaroodi, A. & Bashash, D. The PI3K/Akt/mTOR pathway in lung cancer; oncogenic alterations, therapeutic opportunities, challenges, and a glance at the application of nanoparticles. Transl Oncol 18, 101364, doi:10.1016/j.tranon.2022.101364 (2022).

13 Scrima, M. et al. Signaling networks associated with AKT activation in non-small cell lung cancer (NSCLC): new insights on the role of phosphatydil-inositol-3 kinase. PLoS One 7, e30427, doi:10.1371/journal.pone.0030427 (2012).

14 Gargalionis, A. N., Papavassiliou, K. A. & Papavassiliou, A. G. Implication of mTOR Signaling in NSCLC: Mechanisms and Therapeutic Perspectives. Cells 12, doi:10.3390/cells12152014 (2023).

15 Dobashi, Y., Watanabe, Y., Miwa, C., Suzuki, S. & Koyama, S. Mammalian target of rapamycin: a central node of complex signaling cascades. Int J Clin Exp Pathol 4, 476–495 (2011).

16 Saxton, R. A. & Sabatini, D. M. mTOR Signaling in Growth, Metabolism, and Disease. Cell 169, 361–371, doi:10.1016/j.cell.2017.03.035 (2017).

17 Saxton, R. A., Chantranupong, L., Knockenhauer, K. E., Schwartz, T. U. & Sabatini, D. M. Mechanism of arginine sensing by CASTOR1 upstream of mTORC1. Nature 536, 229–233, doi:10.1038/nature19079 (2016).

18 Li, T. et al. RNF167 activates mTORC1 and promotes tumorigenesis by targeting CASTOR1 for ubiquitination and degradation. Nat Commun 12, 1055, doi:10.1038/s41467-021-21206-3 (2021).

19 Loo, S. K. et al. CASTOR1 phosphorylation predicts poor survival in male patients with KRAS-mutated lung adenocarcinoma. Cell Biosci 14, 127, doi:10.1186/s13578-024-01307-4 (2024).

20 Li, T., Ju, E. & Gao, S. J. Kaposi sarcoma-associated herpesvirus miRNAs suppress CASTOR1-mediated mTORC1 inhibition to promote tumorigenesis. J Clin Invest 129, 3310–3323, doi:10.1172/JCI127166 (2019).

21 Zhou, X. et al. CASTOR1 suppresses the progression of lung adenocarcinoma and predicts poor prognosis. J Cell Biochem 119, 10186–10194, doi:10.1002/jcb.27360 (2018).

22 Huang, M. Y. et al. The DRAP1/DR1 Repressor Complex Increases mTOR Activity to Promote Progression and Confer Everolimus Sensitivity in Triple-Negative Breast Cancer. Cancer Res 84, 2660–2673, doi:10.1158/0008-5472.CAN-23-2781 (2024).

23 Kitai, H. et al. Combined inhibition of KRAS(G12C) and mTORC1 kinase is synergistic in non-small cell lung cancer. Nat Commun 15, 6076, doi:10.1038/s41467-024-50063-z (2024).

24 Luo, Y. D. et al. Mutant Kras and mTOR crosstalk drives hepatocellular carcinoma development via PEG3/STAT3/BEX2 signaling. Theranostics 12, 7903–7919, doi:10.7150/thno.76873 (2022).

25 Awad, M. M. et al. Acquired Resistance to KRAS(G12C) Inhibition in Cancer. N Engl J Med 384, 2382–2393, doi:10.1056/NEJMoa2105281 (2021).

26 Yaeger, R. et al. Molecular Characterization of Acquired Resistance to KRASG12C-EGFR Inhibition in Colorectal Cancer. Cancer Discov 13, 41–55, doi:10.1158/2159-8290.CD-22-0405 (2023).

27 Dobashi, Y. et al. Critical and diverse involvement of Akt/mammalian target of rapamycin signaling in human lung carcinomas. Cancer 115, 107–118, doi:10.1002/cncr.23996 (2009).

28 Balsara, B. R. et al. Frequent activation of AKT in non-small cell lung carcinomas and preneoplastic bronchial lesions. Carcinogenesis 25, 2053–2059, doi:10.1093/carcin/bgh226 (2004).

29 Scheffler, M. et al. PIK3CA mutations in non-small cell lung cancer (NSCLC): genetic heterogeneity, prognostic impact and incidence of prior malignancies. Oncotarget 6, 1315–1326, doi:10.18632/oncotarget.2834 (2015).

30 Dobashi, Y. et al. Paradigm of kinase-driven pathway downstream of epidermal growth factor receptor/Akt in human lung carcinomas. Hum Pathol 42, 214–226, doi:10.1016/j.humpath.2010.05.025 (2011).

31 Hiramatsu, M. et al. Activation status of receptor tyrosine kinase downstream pathways in primary lung adenocarcinoma with reference of KRAS and EGFR mutations. Lung Cancer 70, 94–102, doi:10.1016/j.lungcan.2010.01.001 (2010).

32 Dhillon, T. et al. Overexpression of the mammalian target of rapamycin: a novel biomarker for poor survival in resected early stage non-small cell lung cancer. J Thorac Oncol 5, 314–319, doi:10.1097/JTO.0b013e3181ce6604 (2010).

33 Gately, K. et al. Overexpression of the mammalian target of rapamycin (mTOR) and angioinvasion are poor prognostic factors in early stage NSCLC: a verification study. Lung Cancer 75, 217–222, doi:10.1016/j.lungcan.2011.06.012 (2012).

34 Chantranupong, L. et al. The CASTOR Proteins Are Arginine Sensors for the mTORC1 Pathway. Cell 165, 153–164, doi:10.1016/j.cell.2016.02.035 (2016).

35 Mendoza, M. C., Er, E. E. & Blenis, J. The Ras-ERK and PI3K-mTOR pathways: cross-talk and compensation. Trends Biochem Sci 36, 320–328, doi:10.1016/j.tibs.2011.03.006 (2011).

36 Judd, J. et al. Characterization of KRAS Mutation Subtypes in Non-small Cell Lung Cancer. Mol Cancer Ther 20, 2577–2584, doi:10.1158/1535-7163.MCT-21-0201 (2021).

37 Wood, K., Hensing, T., Malik, R. & Salgia, R. Prognostic and Predictive Value in KRAS in Non-Small-Cell Lung Cancer: A Review. JAMA Oncol 2, 805–812, doi:10.1001/jamaoncol.2016.0405 (2016).

38 Bakhoum, S. F. & Cantley, L. C. The Multifaceted Role of Chromosomal Instability in Cancer and Its Microenvironment. Cell 174, 1347–1360, doi:10.1016/j.cell.2018.08.027 (2018).

39 Megyesfalvi, Z. et al. Clinical insights into small cell lung cancer: Tumor heterogeneity, diagnosis, therapy, and future directions. CA Cancer J Clin 73, 620–652, doi:10.3322/caac.21785 (2023).

40 Ma, L., Chen, Z., Erdjument-Bromage, H., Tempst, P. & Pandolfi, P. P. Phosphorylation and functional inactivation of TSC2 by Erk implications for tuberous sclerosis and cancer pathogenesis. Cell 121, 179–193, doi:10.1016/j.cell.2005.02.031 (2005).

41 Castellano, E. & Downward, J. RAS Interaction with PI3K: More Than Just Another Effector Pathway. Genes Cancer 2, 261–274, doi:10.1177/1947601911408079 (2011).

42 She, Q. B. et al. 4E-BP1 is a key effector of the oncogenic activation of the AKT and ERK signaling pathways that integrates their function in tumors. Cancer Cell 18, 39–51, doi:10.1016/j.ccr.2010.05.023 (2010).

43 Carracedo, A. et al. Inhibition of mTORC1 leads to MAPK pathway activation through a PI3K-dependent feedback loop in human cancer. J Clin Invest 118, 3065–3074, doi:10.1172/JCI34739 (2008).

44 Turke, A. B. et al. MEK inhibition leads to PI3K/AKT activation by relieving a negative feedback on ERBB receptors. Cancer Res 72, 3228–3237, doi:10.1158/0008-5472.CAN-11-3747 (2012).

45 Schram, A. M. et al. A phase Ib dose-escalation and expansion study of the oral MEK inhibitor pimasertib and PI3K/MTOR inhibitor voxtalisib in patients with advanced solid tumours. Br J Cancer 119, 1471–1476, doi:10.1038/s41416-018-0322-4 (2018).

46 Engelman, J. A. et al. Effective use of PI3K and MEK inhibitors to treat mutant Kras G12D and PIK3CA H1047R murine lung cancers. Nat Med 14, 1351–1356, doi:10.1038/nm.1890 (2008).

47 DuPage, M., Dooley, A. L. & Jacks, T. Conditional mouse lung cancer models using adenoviral or lentiviral delivery of Cre recombinase. Nat Protoc 4, 1064–1072, doi:10.1038/nprot.2009.95 (2009).

48 Hai, J. et al. Generation of Genetically Engineered Mouse Lung Organoid Models for Squamous Cell Lung Cancers Allows for the Study of Combinatorial Immunotherapy. Clin Cancer Res 26, 3431–3442, doi:10.1158/1078-0432.CCR-19-1627 (2020).

49 Ludwig, J. et al. An easy-to-perform protocol for culturing primary murine lung tumor cells as organoids. Ann Anat 255, 152298, doi:10.1016/j.aanat.2024.152298 (2024).

50 Ramos da Silva, S., et al. Broad Severe Acute Respiratory Syndrome Coronavirus 2 Cell Tropism and Immunopathology in Lung Tissues From Fatal Coronavirus Disease 2019. J Infect Dis 223, 1842–1854, doi:10.1093/infdis/jiab195 (2021).

