## Supplementary figures and images for "*CASTOR1*: A Novel Tumor Suppressor Linking mTORC1 and KRAS Pathways in Tumorigenesis and Resistance to KRAS-Targeted Therapies in Non-Small Cell Lung Cancer"

### Supplemental Materials

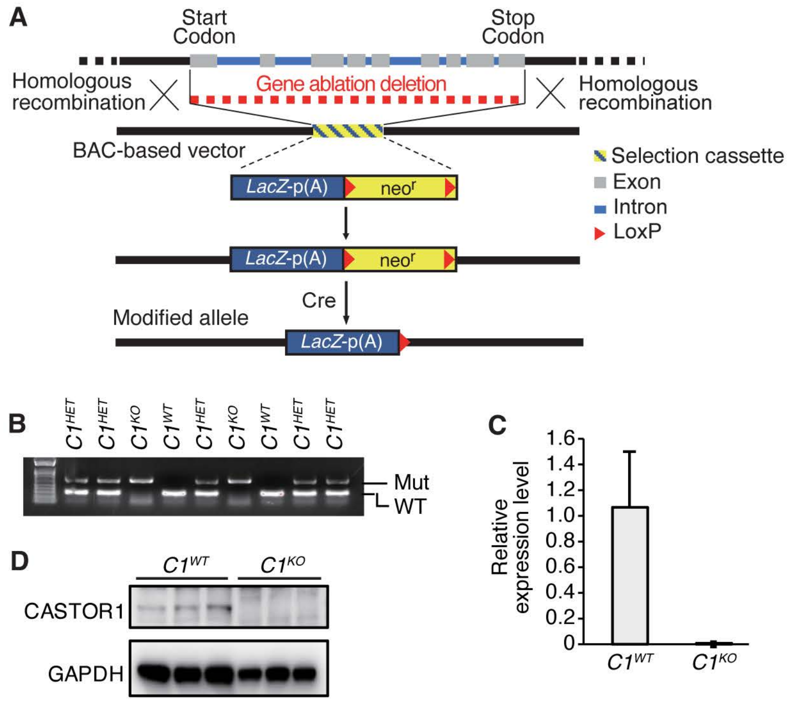

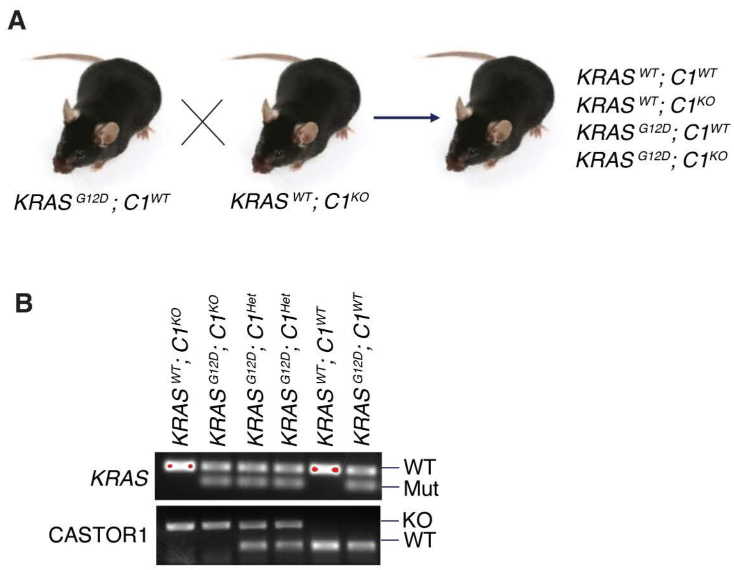

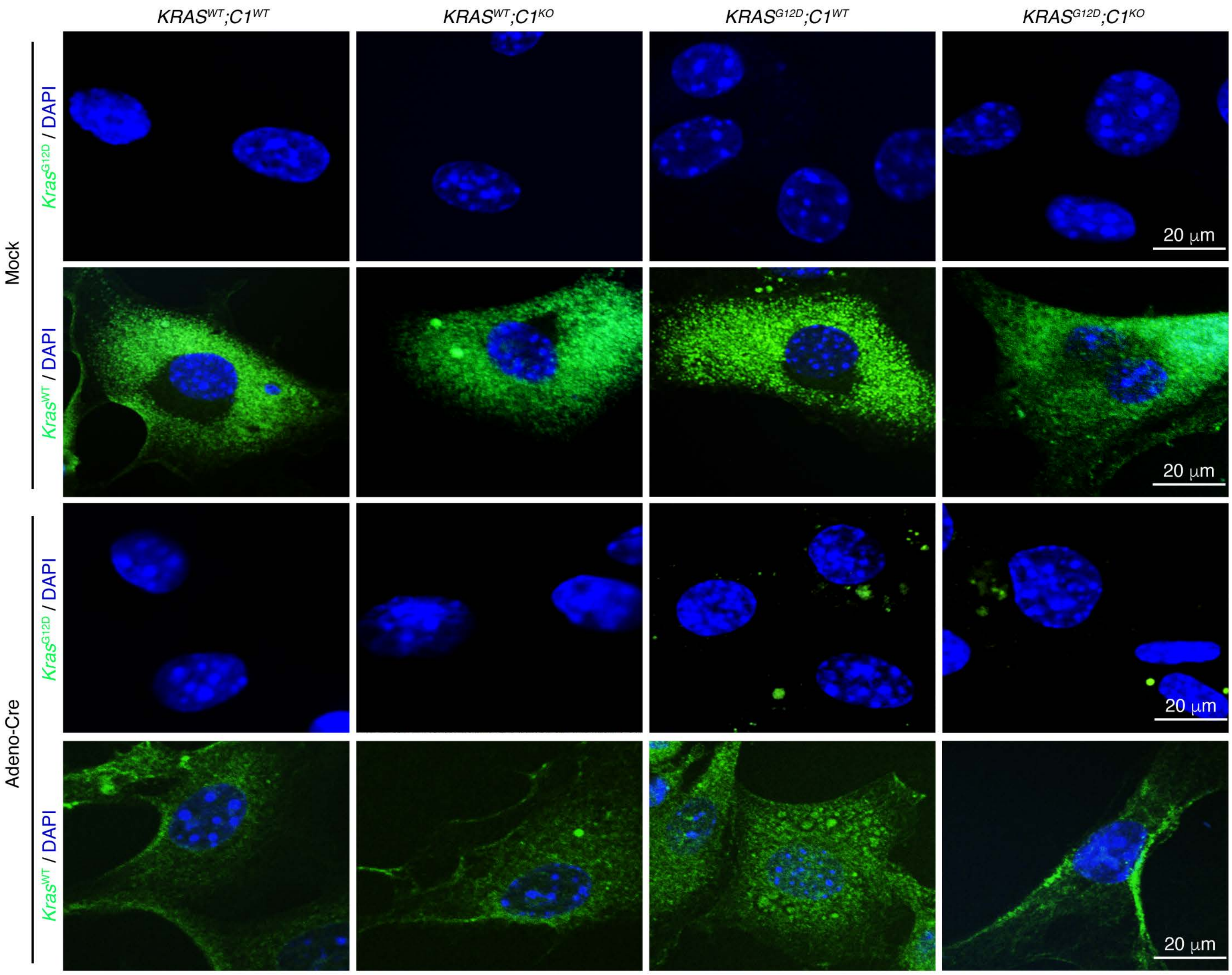

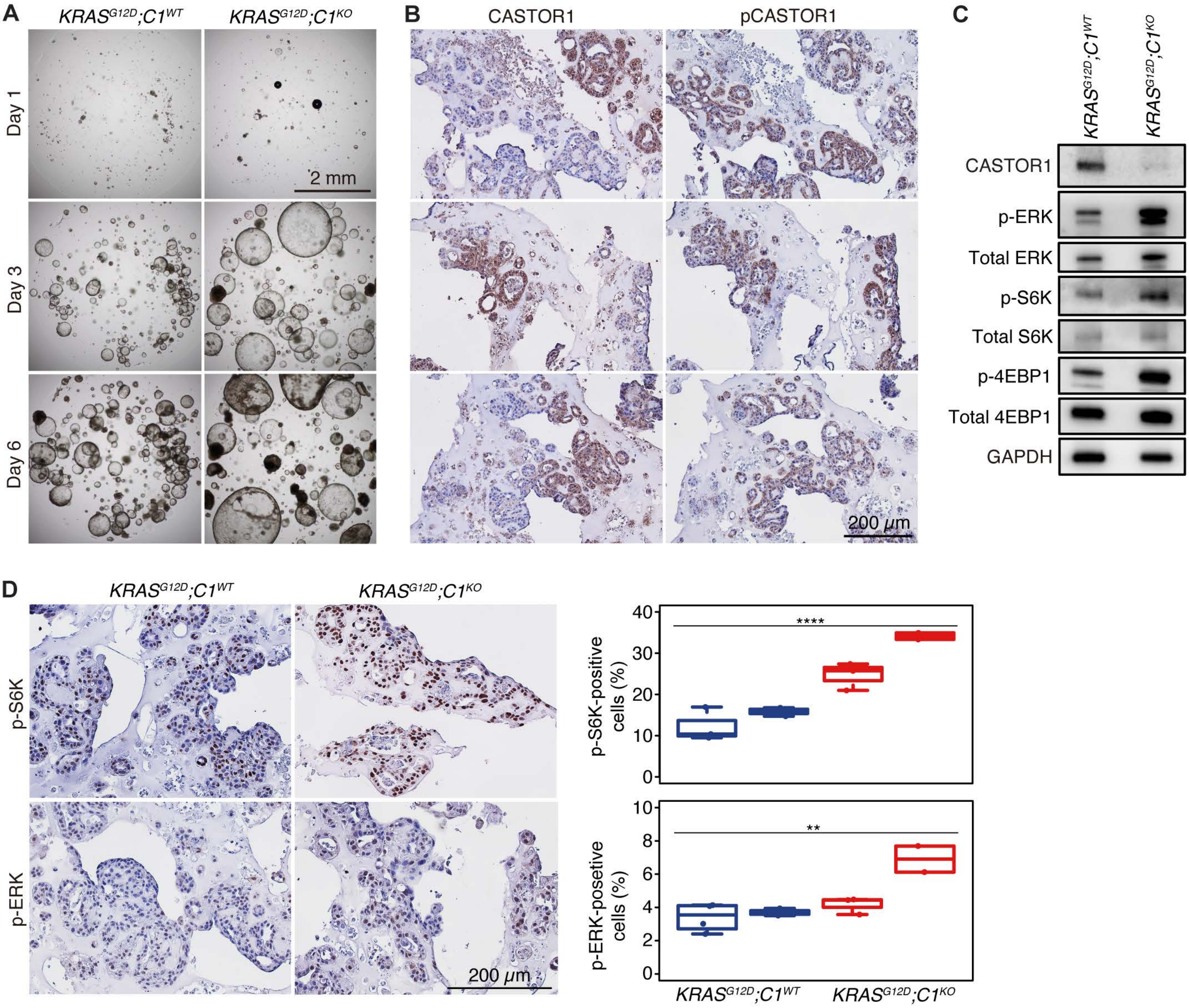
